# Model-based prediction and ascription of deforestation risk within commodity sourcing domains: Improving traceability in the palm oil supply chain

**DOI:** 10.1101/2024.05.01.589041

**Authors:** Henry B. Glick, Judith M. Ament, Jorn S. Dallinga, Juan Torres-Batlló, Megha Verma, Nicholas Clinton, Andrew Wilcox

## Abstract

Palm oil accounts for approximately 50% of global vegetable oil production, and trends in consumption have driven large-scale expansion of oil palm (*Elaeis guineensis*) plantations in Southeast Asia. This expansion has led to deforestation and other socio-environmental concerns that challenge consumer goods companies to meet no deforestation and sustainability commitments. In support of these commitments and supply chain traceability, we seek to improve on the current industry standard sourcing model for ascribing social and environmental risks to particular actors. Using passive geolocational traceability data (***n*** = 3,355,437 cellular pings) and machine learning models, we evaluate the industry standard sourcing model, and we predict with high accuracy the undisclosed sourcing domains for 1,570 Indonesian and Malaysian palm oil mills on the Universal Mill List (as of November 2021). In combination with the World Wide Fund for Nature – Netherlands’ Forest Foresight, we use our predicted sourcing domains to provide an illustrative example of the proportional allocation of future deforestation, carbon loss, and biodiversity risk to relevant actors, permitting targeted outreach, contract negotiation, and mitigation of large-scale resource degradation. This example depends on a subset of model predictions in the absence of disclosed traceability data. The utilization of additional predictions or disclosed traceability data would influence and improve the results.

## Main

Global consumption of palm oil and palm oil derivatives (hereafter, palm oil) has more than doubled in the past two decades, accounting for nearly 50% of global vegetable oil production [1, 2]. This trend has driven large-scale expansion of palm plantations [3, 4], particularly in Southeast Asia (Indonesia and Malaysia), where production is an order of magnitude greater than other palm oil producing regions [1, 5]. Where commodity production has emerged as the dominant driver of deforestation globally [3, 6, 7], consumer goods companies and those utilizing deforestation risk-tied commodities, have elected, or been pressured [7, 8], to mitigate continued deforestation through supply chain commitments. By 2017 nearly 450 companies had implemented 760 corporate sustainability commitments to more “sustainable” sourcing [9, 10].

Despite these commitments, the complexity of the palm oil supply chain presents a major barrier to their widespread success [10–12]. Although the rate of deforestation across Indonesia and Malaysia has slowed markedly in recent years (an average year-on-year reduction in forest loss of 7.2% from 2015-2021, or a 51% reduction overall [13], the ubiquity of oil palm production, high yield [14], access to local and/or global markets, and hopes of economic prosperity continue to drive industry growth at both industrial and smallholder scales [11]. In turn, deforestation [3], land use conversion [4], and degradation of critical landscapes [5] continue, spurring downstream issues associated with biodiversity loss [15], resource-based conflict [16, 17], and human rights abuses [18–20]. While sustainability certifications (e.g., Roundtable on Sustainable Palm Oil; [21, 22]; or Indonesia Initiative for Sustainable Palm Oil [23, 24]) and top down policy decisions from national governments intend to suppress deforestation, land use conversion, and social conflict, sustainability commitments require that consumer goods companies manage their supply chains through targeted risk assessment. In the present paper, we are concerned with the manner in which risk is ascribed to relevant actors.

We define risk to be the perceived risk, on behalf internal and external stakeholders, of violating company-wide sustainability commitments, whatever those may be. Implicit in this definition is the notion that risk is subjective. This is highlighted by the vast assortment of corporate sustainability commitments [9, 10], each of which perceives risk through a slightly different lens. In recent years, the “no deforestation” commitments of palm oil consumer goods companies have evolved into more holistic risk reduction targets (e.g., NDPE, or No Deforestation, Peat conversion, and human Exploitation; [25, 26], https://palmoilcollaborationgroup.net/), including Net Zero [27, 28] and Scope 3 [29, 30] emissions commitments associated with land use change. Yet, while acknowledging the positive trends achieved in palm oil production, when we consider all sources of loss, since 2000, nearly 7% of total primary tropical forest cover has been lost at an average pace of 3.30 million hectares per year [13], driven largely by commercial agriculture [6, 31]. Year 2021 saw the loss of 25.3 million hectares of tree cover globally, 3.75 million hectares of which consisted of tropical primary rainforests, responsible for 2.46 Gt of carbon dioxide-equivalent emissions (Indonesia and Malaysia represent 7.3% of this loss and 8.6% of these emissions) [13].

In the context of consumer goods companies, risk assessment generally depends on two mutualistic components: (i) consensus among deciding parties on what constitutes risk, and (ii) ascription of risky or non-risky activities and behaviors to the appropriate actors. In the case of palm oil production, the actors are primarily independent traders, mills, and refineries that are in-network for a given consumer goods company. With knowledge of which actors are leading to elevated risk, consumer goods companies can identify those actors for discussion, remediatory action, penalty, or exclusion from the supply chain, ultimately enabling a change in behavior through social or economic pressures. In recent years, consumer goods companies have relied on a number of spatially explicit risk assessment tools (models) to define risk over select geographic domains. In the case of the palm oil supply chain, examples include [32–37], which intend to quantify risk of either deforestation (e.g., [37]) or NDPE more broadly (e.g., [33, 38]).

Even with spatially explicit encapsulations of risk, we still face a fundamental challenge in ascription of that risk. For years, NGOs and multinational companies in the palm industry have relied on a *de facto* 50 km radial sourcing model, in which a circular region 100 km in diameter is circumscribed around a facility [32, 38–40]. After accounting for known concessions and/or plantations, the entirety of the land within this circular region, and by extension, any risky or non-risky activity within this circular region, is then ascribed to the focal facility. This approach was borne out of pragmatism and limitations in data availability. A 50 km radial distance reflects a pairing of local infrastructural limitations and estimated travel times, providing some form of monitoring over more than 700,000 ha, while maintaining constraints on the quality of palm fresh fruit bunches (FFB). To minimize the degrading effects of free fatty acid development, a FFB needs to be processed no more than 8-48 hours after harvest [41, 42]; a number of companies target an upper limit of 24 hours [43–45]. With respect to data limitations, palm oil suppliers rarely elect to disclose farm- or plantation-level sourcing locations in an effort to maintain a competitive edge in the local commodity marketplace, limit pressure from external stakeholders related to environmental, social, and/or governance (ESG) practices (e.g., [8, 46–48]), and to adhere to government policies (memorandums of understanding) that seek to limit the distribution of potentially sensitive information without proper control and monitoring [49–53]. Despite its prevalence, the radial model reflects a naive approach and lacks quantitative support beyond a crude estimation of the transportation duration tolerated by oil palm FFB.

Here, we present a two-part study intended to enhance the state of spatially explicit commodity-driven risk assessment, and in particular, traceability in the palm oil supply chain. In the first part, we use non-parametric machine learning models and anonymized cellular traceability data linked to palm mills, to: (a) evaluate, for the first time, the industry standard radial sourcing model, and (b) predict with high accuracy the palm oil sourcing domains for each of 1,570 mills on the Universal Mill List [54] and in Unilever’s previously disclosed supply chain. The industry standard radial sourcing model neither draws on the available sourcing context (e.g., palm plantations, human settlement patterns) nor offers any measure of reliability. As a discretized in-out sourcing model, it also fails to address fundamental issues of overlapping sourcing domains, leading to redundant ascription of social or environmental concerns. Here we present an alternative approach focused on relative ownership and the proportional allocation of risky behaviors to relevant actors.

In the second part, we integrate our sourcing domain predictions with the World Wide Fund for Nature – Netherlands’ Forest Foresight [55, 56] — a machine learning-based early warning system, providing near-term predictions of deforestation events. By integrating Forest Foresight deforestation predictions with refined sourcing domains, we are able to ascribe predicted future deforestation events, carbon loss, and biodiversity threat abatement metrics to relevant actors. The combination of these analytical methods provides a framework for addressing risk — deforestation or otherwise — in a manner not previously explored. Addressing violations to sustainability commitments need not be solely responsive to past events, and ascription of risky behavior to relevant actors permits targeted outreach, ultimately enhancing commodity supply chains for the benefit of suppliers, consumers, and the planet. Though our focus here is on deforestation and the palm oil supply chain, the methods we present are crop, risk, and geography agnostic.

## Results

### Example mills

We selected three palm processing mills to serve as examples in presenting mill-level modeling results (Table 1). These mills are located in Seruyan and Kotawaringin Barat (West Kotawaringin) regencies in Central Kalimantan, Indonesia, an area containing 63 other palm mills within 50 km. These facilities were selected because (a) they are previously disclosed suppliers in Unilever’s palm oil supply chain, (b) there is coincident Forest Foresight data available for their locations, and (c) they represent varied certification status, supplier competition, geographic placement (coastal versus inland), and training data sample sizes. We emphasize that these mills serve only to demonstrate how our methods and data might be used, and statements made regarding mill and/or supplier responsibility (a) are informed by model-based predictions that do not necessarily reflect an objective truth; and (b) reflect the application of our methods to a limited set of data. Inclusion of fewer or additional example mills, or of disclosed traceability data, would produce an alternate set of results.

**Table 1:**
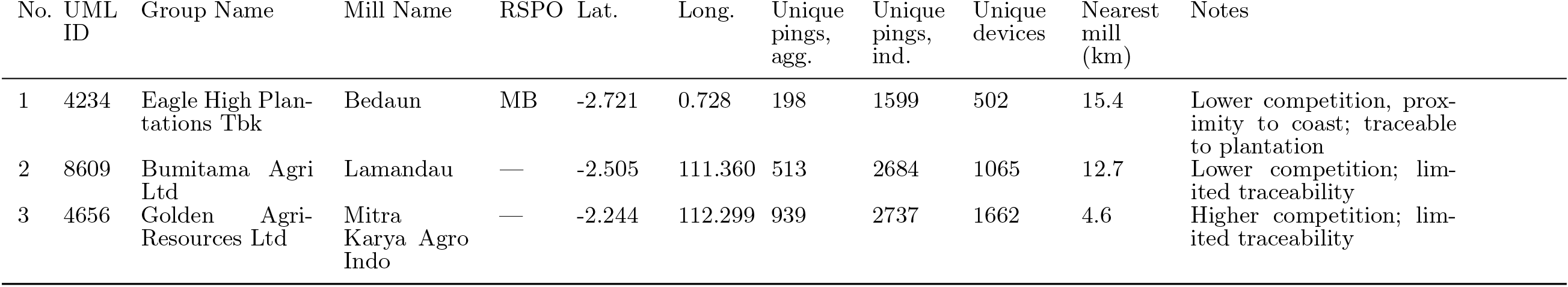
Example mills used to highlight mill-level modeling results. Modeling was based on anonymized aggregated cellular pings (Unique pings, agg.). UML ID = Universal Mill List identifier. All UML ID values are preceded with PO100000; RSPO = Round Table on Sustainable Palm Oil certification status (MB = mass balance); agg. = aggregated; ind. = individual.

### Mill-level results

As applied to each of the example mills in Table 1, Figure 1 shows the accuracy assessment metrics for four predictive modeling approaches that seek to improve on the 50 km radial approach. These include three individual approaches — maximum entropy (MaxEnt;) [57–60]), down-sampled random forest (RF; [61, 62], and down-sampled gradient boosted regression trees (BRT; [63–65]) — and an ensemble of these three [66, 67]. Across sample sizes, we see that the receiver operating characteristic curves (ROC; [68]), the precision-recall gain curves (PRG; [69]), and the true skill statistic curves (TSS; [70]) far exceed the “no skill” models, and generally approximate their idealized shapes in which the curves reach the upper lefthand and upper righthand corners of each plot. While values for area under the curve (AUC) can be difficult to compare across applications, our individually reported AUC values generally reflect “outstanding discrimination” [71]. In tandem with the maximized precision-recall gain-based F_1_ accuracy (PRGF_1_) [69, 72, 73] and TSS (equivalent to Youden’s J, per [74]), these models demonstrate strong predictive strength. With respect to our individual models, by measures of individual model AUC, our RF models (column b) performed least well in the majority of cases, while our BRT (column c) performed best in the majority of cases. When considering individual model binary classification metrics (PRGF_1_, TSS/Youden’s J), our MaxEnt model performed least well in the majority of cases, succeeded by RF and BRT. Like Valavi *et al*. (2021a), our ensemble of these three models (column d) demonstrated the best performance on a mill-by-mill basis, though the gains in accuracy may not outweigh the computational costs.

**Fig. 1:**
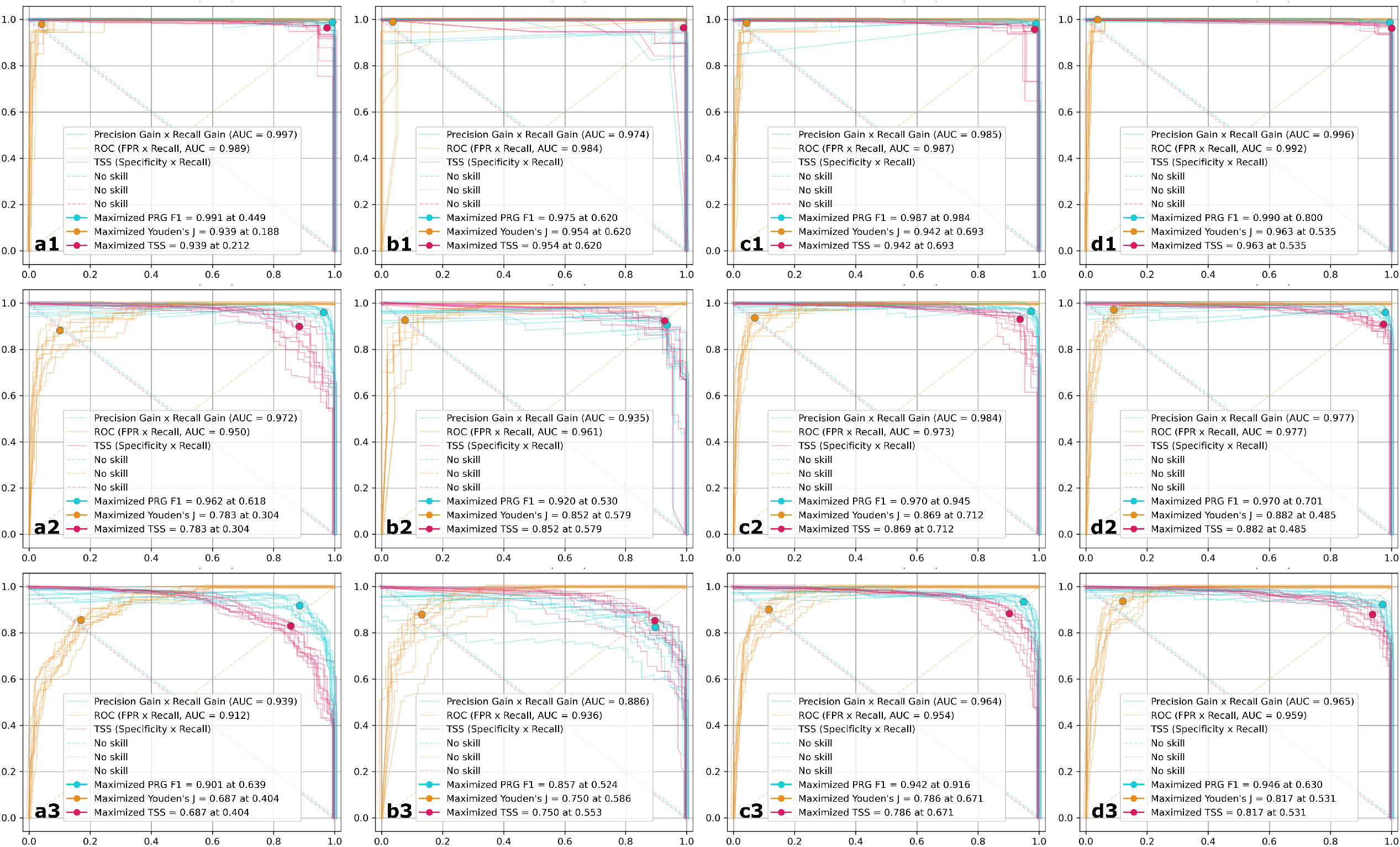
Threshold-independent and threshold-dependent accuracy assessment measures produced for each model (rows) at three sample three mills (columns). a = MaxEnt, b = down-sampled random forests, c = down-sampled gradient boosted regression trees, d = mean ensemble. Mills number 1, 2, and 3 per Table 1. The axis labels for each curve are unique and are noted parenthetically in each legend. Each curve-based metric shows 10 curves derived from 10-fold cross validation. Plotted color-coded points represent the average values of the 10 maximized threshold-dependent metrics (PRGF_1_, Younden’s J, and TSS). As averages, these values are informative only, and do not actually exist in any one set of predictions. For the color-coded points, the reported values are the fold-level maximum values for each respective metric. Mathematically, Youden’s J and TSS are maximized at the same thresholds, but their curves are visualized on different axes. Curve coordinates have been jittered randomly (*±*0.8%) for visual clarity.

Figure 2 provides examples of predicted sourcing probability surfaces for the mills detailed in Table 1 and Fig. 1. The reader will note the spatial variability in probability values, driven by measures of travel cost and mill-to-mill competition. The restricted predictions of mill 1 are a function of spatially consolidated training data, in turn suggesting that this particular mill may source most of its palm from a limited area. Figure S5 captures the classified (binary, sourcing versus non-sourcing domain) versions of each surface shown in Fig. 2. Classification was conducted with the average of thresholds that optimized PRGF_1_ accuracy. Information is lost in the classification of the continuous surfaces, but doing so mirrors the current industry standard model and generally represents the data structures used by NGOs and consumer goods companies. In Fig. S5, the industry standard model (red circle) greatly exceeds the predicted sourcing domains in nearly all cases. This indicates that our preicted sourcing domains could ultimately improve supplier-level outlook by helping them to better focus their resources.

**Fig. 2:**
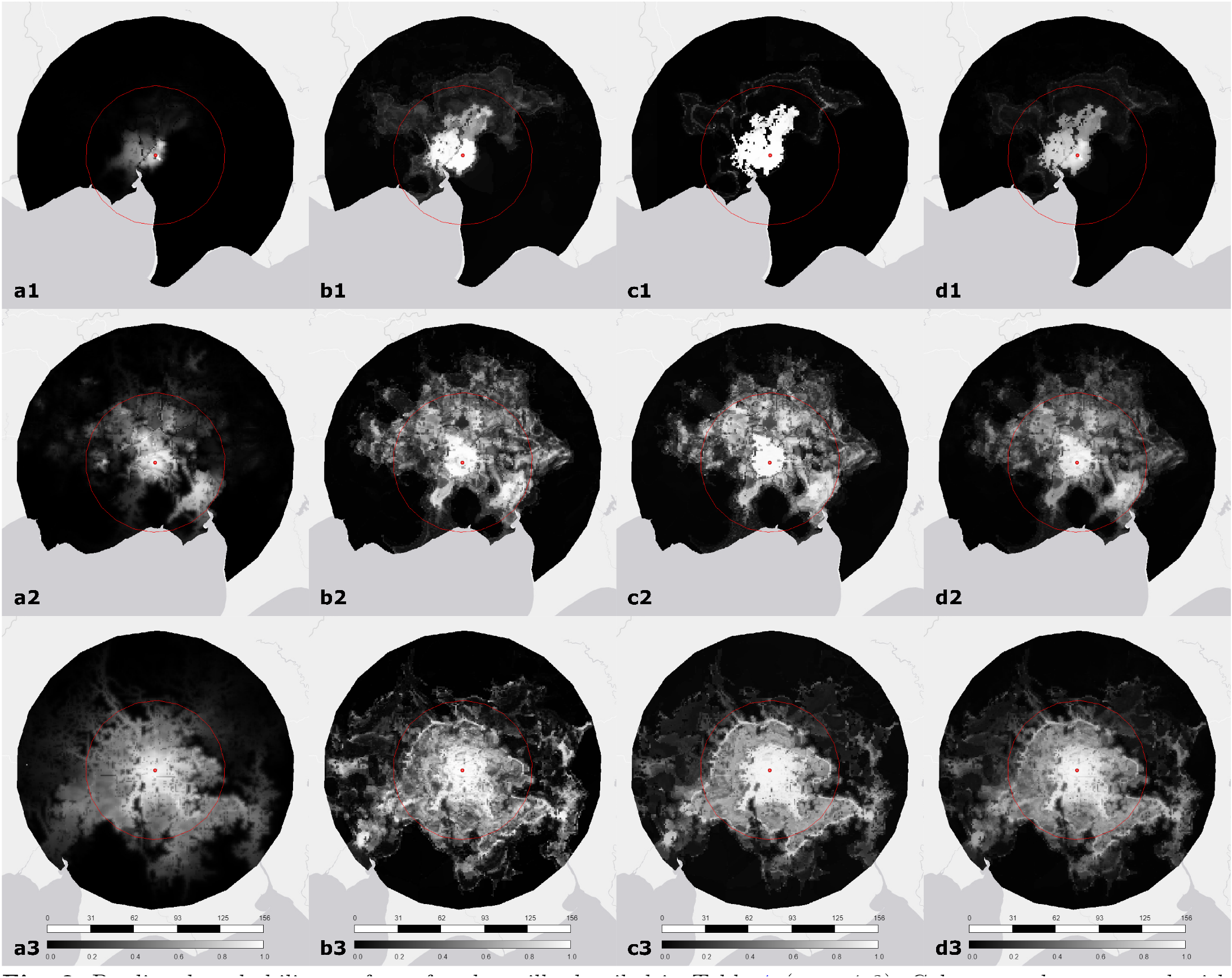
Predicted probability surfaces for the mills detailed in Table 1 (rows 1-3). Columns a-d correspond with MaxEnt, down-sampled random forests, down-sampled gradient boosted regression trees, and ensemble models, respectively. Red circular region corresponds with the current industry standard 50 km radial sourcing model. Red point marks mill location. Legend: upper bar is scale, in km; lower bar is probability.

### Regional results

Figure 3 highlights the core findings of this study, enabling cross-mill evaluation of our models against one another, and against the industry standard. The thresholds used to maximize the TSS and PRGF_1_ accuracy (left pane) are less unified than anticipated. In the context of unbalanced data with pseudo-absences and a preference for reduction of false positives over false negatives, we would generally seek to optimize TSS thresholds ≤ 0.5 and PRGF_1_ thresholds of ≈ 0.5 (cf [75] Fig. 1). Our RF and BRT thresholds for TSS, and our BRT and ensemble thresholds for PRGF_1_, are higher than expected. For both the TSS and PRGF_1_ thresholds, all metrics were significantly different from one another. With respect to the threshold-dependent metrics (TSS, PRGF_1_, Kappa; [76]), the rank order of models for maximum TSS and maximum PRGF_1_ metrics generally mirrors those shown in Fig. 1. The maximum PRGF_1_ values indicate that the RF models are of lesser strength, though the individual models all yield high average maximum PRGF_1_, with less variable distributions than those of the maximum TSS values.

**Fig. 3:**
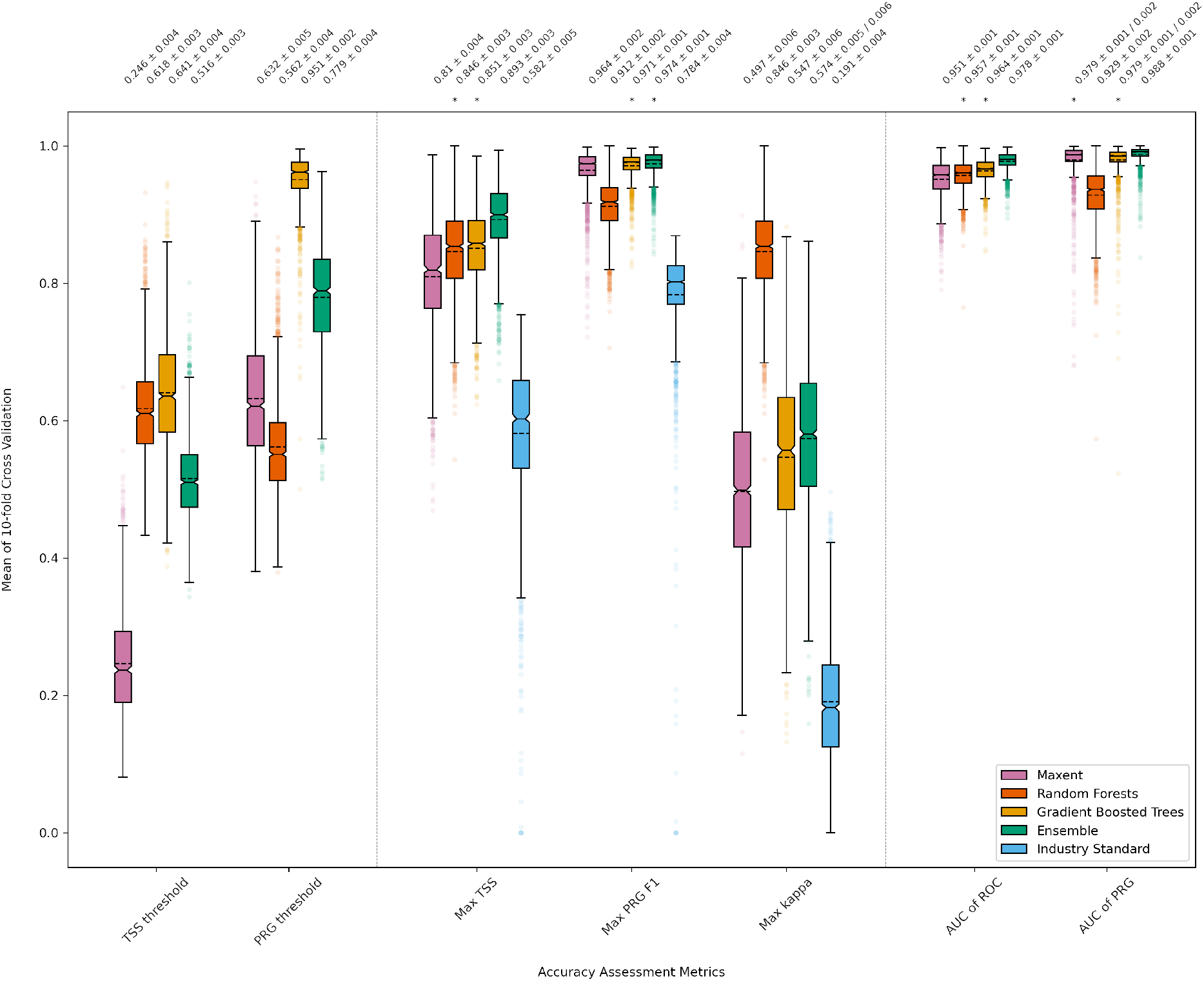
Summary measures for all mill-level models (1,570 mills *×* 10-fold cross validation *×* 5 models = 78,500 models). Each box represents 1,570 10-fold cross validation means from mill-level models, with median (solid) and mean (dashed) lines. Left pane = the thresholds at which TSS and PRG were maximized. Middle pane = the maximum of each respective figure across cross validation folds. Right pane = the area under the receiver operating characteristic curve and precision-recall gain curve. Models with an asterisk are not significantly different (*α* = 0.05) for that particular metric. Values in upper margin are means *±* margins of error from 95% bootstrapped confidence intervals.

We have included Cohen’s kappa [76] in Fig. 3, but with its inherent limitations (see Supplementary Information § Tuning subset, variable selection, and accuracy assessment), differences should be considered carefully. Across TSS, PRGF_1_, and kappa, we see overwhelming evidence that the industry standard model performs significantly worse than our models. Allowing that kappa is sensitive to prevalence, the distribution of industry standard kappa values reflects a status quo that is hardly better than random chance. With respect to the AUROC and AUPRG shown in Fig. 3, we see that all models perform excellently, in some cases nearly maximizing the theoretical limits of 100% AUC. On average, our ensembles perform significantly better than any of the individual models, with the worst mill-level ensemble model capturing 89.4% and 83.8% of AUROC and AUPRG, respectively. There is at least one mill-level model for each model type that achieved, under 10-fold cross validation, an AUROC and AUPRG of approximately 1.

### Ascription of deforestation risk, carbon emissions, and biodiversity loss

#### Example ascription of forest and carbon loss by hot zone

Figure 4 depicts a single Forest Foresight “hot zone” – a down-sampled region 480 m *×* 480 m, tagged with the areal proportion predicted to be at risk of deforestation, as derived from predictions made at 15 m *×* 15 m [55] (see § Methods) – from July 2021 (red square, coordinates [111.57062634879999, -2.6929287232], index 78026), following closely on the temporal span of our traceability data (see § Methods). The hot zone was predicted to contain deforestation in the six month period following July 2021. Fig. 4a shows the state of the hot zone prior to the date predictions were made, while Fig. 4b shows the state of the hot zone following the six month prediction window, with clear signs of deforestation. The reader will note the exposed ground and two-tone striations in Fig. 4b, which reflect windrowing of cut trees and slash in preparation for burning – a common practice in the installation of palm plantations. Based on real-time alerting, deforestation in and around the hot zone took place from September through November of 2021, falling squarely in the six month prediction window Forest Foresight sought to capture.

**Fig. 4:**
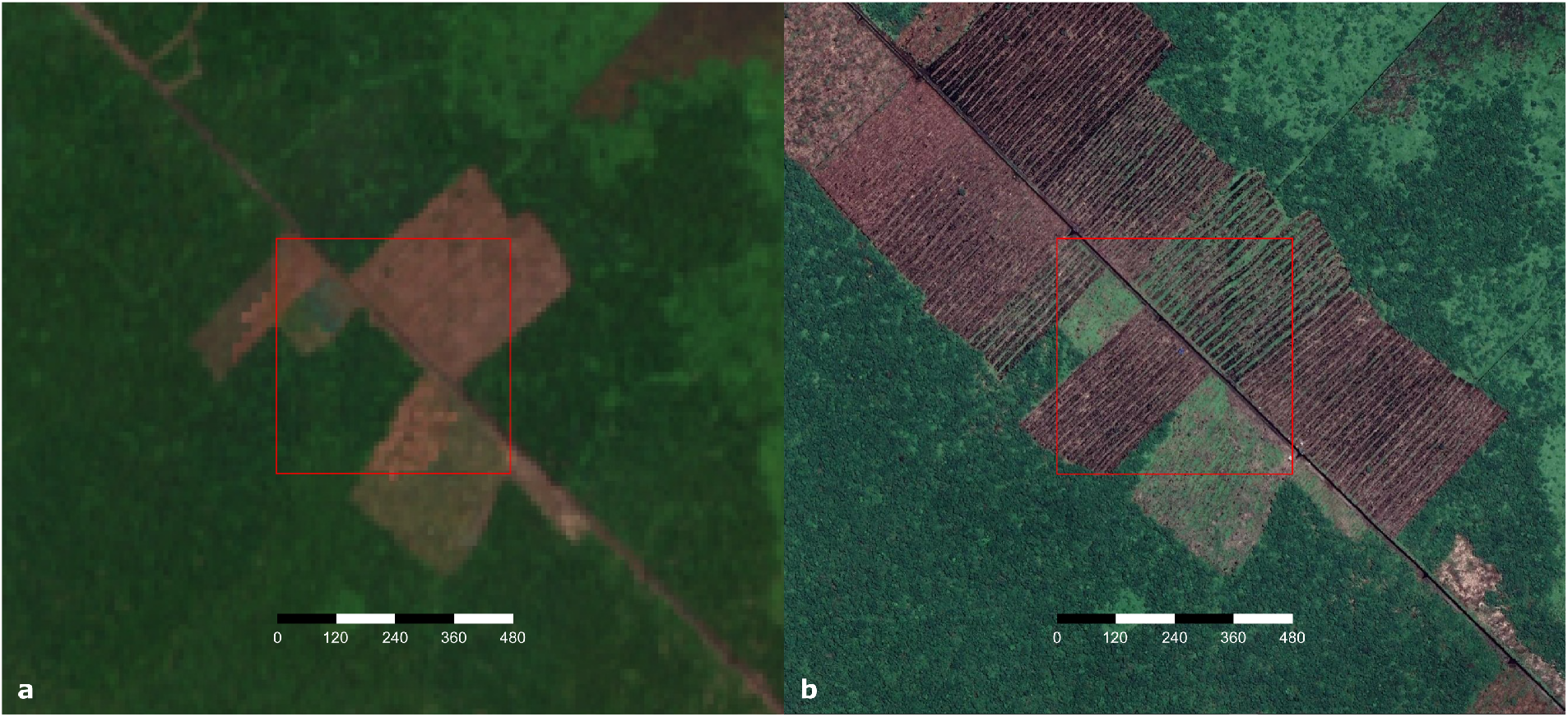
Forest Foresight hot zone index 78026 (red square) in Central Kalimantan. Sentinel-2 true color composite from imagery captured between January 1 and June 30, 2021 (left; 10 m spatial resolution), and high resolution true color Google basemap imagery captured in year 2022 (right; *<*1 m spatial resolution; © CNES, AirBus, Maxar). Year 2022 basemap imagery shows recent deforestation and windrowing of trees and slash in preparation for burning (visible as dark rows in some plots), following Forest Foresight predictions from July 2021. Scale bar in meters.

Of the 230,827 square meters (per location-appropriate equal-area projection) in the hot zone, 43.8%, or 10.11 ha, was predicted by Forest Foresight to be at risk of near-term deforestation. The total above-ground forest carbon (C) in the hot zone was estimated to be 1356.11 MgC as of mid-year 2020. Using the three mills detailed in Table 1, we present the mill-wise allocation vector and proportional forecasted carbon loss for this hot zone in Table 2A. With the acknowledged limitations of applying our approach to the example mills, Table 2A illustrates that for this particular hot zone, the Mitra Karya Agro Indo mill is predicted to hold the majority of the responsibility for the forecasted (and realized) deforestation. Fig. 4b suggests that there has been, or will soon be, a direct loss of carbon storage from burning. The reader will note that the carbon density within this hot zone, as derived from the allocated total carbon and allocated total area within the hot zone, is relatively low (58.6 MgC/ha) compared to the Indonesian national average (142-158 MgC/ha, per [77]). Forest Foresight predictions include an estimated areal proportion of each hot zone at risk of deforestation (see Table 2). In this particular example, where we can clearly see the near total loss of above-ground forest carbon outside of the remaining in-tact forested area (Fig. 4b), we can make use of this estimate of at-risk forested area to derive a carbon density of 133.7-136.5 MgC/ha (allocated total C / allocated forest at risk), as would be more typical of this region. This approach is only possible because of the close inspection of the hot zone. Although Forest Foresight hot zones may contain, or otherwise fall within close proximity to existing deforestation, the predictions do not make direct claims about the non-at-risk portions of each hot zone. In the current data product, deforestation may be predicted in hot zones that are otherwise fully forested or which contain variable above-ground carbon density.

**Table 2:**
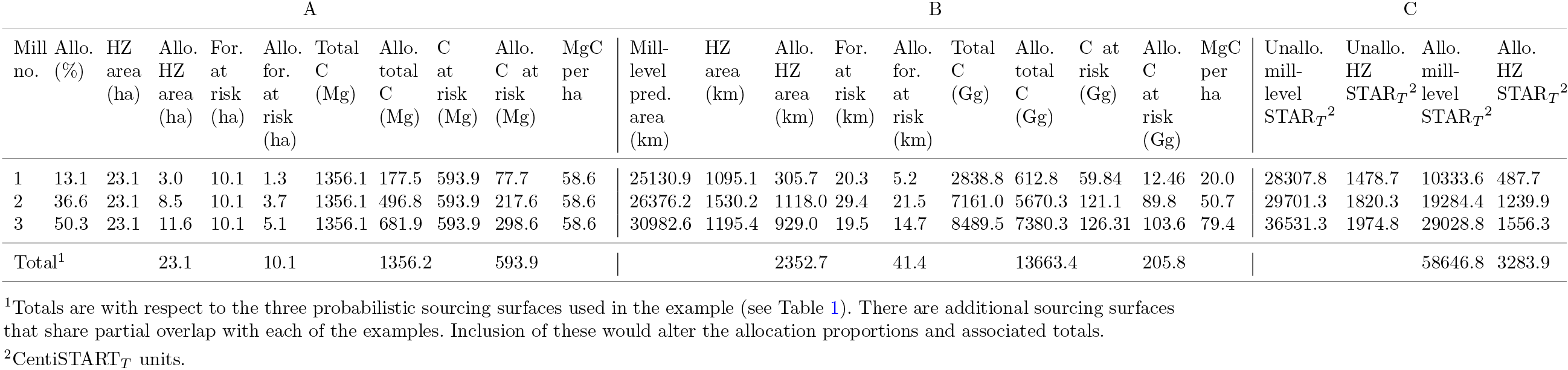
Example ascription of predicted deforestation, aboveground forest carbon (C) loss, and biodiversity threat using our ensemble model. (A) Deforestation and carbon loss metrics based on Forest Foresight hot zone index 78026. (B) Deforestation and carbon loss metrics based on all Forest Foresight hot zones (July 2021) falling within the prediction domains of the mills shown in Table 1. (C) Like (B) but for Species Threat Abatement Metrics. For. = forest, Allo. = allocation/allocated, Unallo. = unallocated, HZ = Forest Foresight hot zone, pred. = prediction. MgC/ha computed from allocated total MgC/allocated total hot zone area.

### Mill-level ascription of forest, carbon loss, and biodiversity risk

Table 2B and 2C present summarizing measures for all Forest Foresight hot zones encapsulated by the maximal prediction domains for each of the three example mills detailed in Table 1. Here we see that while comparable in total hot zone area, the Bedaun mill (no. 1) is predicted to hold far less responsibility for the landscape-level forecasted deforestation, carbon loss, and biodiversity risk than do the other two mills. From the perspective of a consumer goods company, a prioritized effort to tackle violations to NDPE commitments would place energy on purchasing associated with the Mitra Karya Agro Indo mill (no. 3), followed by the Lamandau mill (no. 2), and then Bedaun (no. 1). We emphasize again, that these statements are informed by our models and available data, and do not necessarily reflect an objective truth. There are additional sourcing surfaces that share partial overlap with each of the examples. Inclusion of these would alter the allocation proportions and associated totals.

The total hot zone-level allocations of predicted forest loss (ha), carbon loss (MgC), and biodiversity risk (centiSTAR_*T*_, or Species Threat Abatement units, per [78]; see 9Methods) are not particularly large for any of the example mills. The reason for this lies in the landscape structure around existing palm oil processing facilities, which are generally situated in vast matrices of industrial and small-holder palm plantations (Fig. 5). However, when looking across all of Indonesian Borneo, there are 380 mills for which a single set of predictions from July 2021 yields approximately 84,502 hectares of predicted forest loss, 418,661 Gg of predicted carbon loss, and 77,091 centiSTART units of predicted biodiversity risk for the subsequent 6-month prediction window. Here we stress that the predicted forest (and carbon) loss does not discriminate between palm and non-palm outcomes, and instead reflects *any* predicted forest loss. In Southeast Asia, Forest Foresight data is not currently available for mills outside Indonesian Borneo. However, if we allow that deforestation rates on Borneo loosely approximate those in other portions of the region (acknowledging that rates on Java and Sumatra are likely to be lower due to lower remaining forest cover), and multiply the above predictions by a naive scaling factor to estimate the region-wide figures (1,676 regional mills / 380 mills in Kalimantan, covered by Forest Foresight = 4.41 approximate scaling factor), these estimates become: 372,699 ha of predicted forest loss, 1,846,513,486 Mg of predicted carbon loss (1,846.5 TgC across all area within each hot zone; 36.6 TgC if considering only the Forest Foresight estimated areal proportion of each hot zone at risk), and 77,091 centiSTAR_*T*_ units of predicted biodiversity risk. These figures are for a given month’s predictions across the 1,676 Indonesian and Malaysian palm oil mills in Unilever’s previously disclosed supply chain (as of November 2021) and appearing on the Universal Mill List. If we allow that Forest Foresight produces updated (but not mutually exclusive) predictions at monthly intervals (forecast over six-month prediction windows), with 70-80% user’s accuracy and a 50% detection rate [55], these figures begin to demonstrate the potential and broad applicability of predictive models in supply chain traceability and monitoring of resource degradation.

**Fig. 5:**
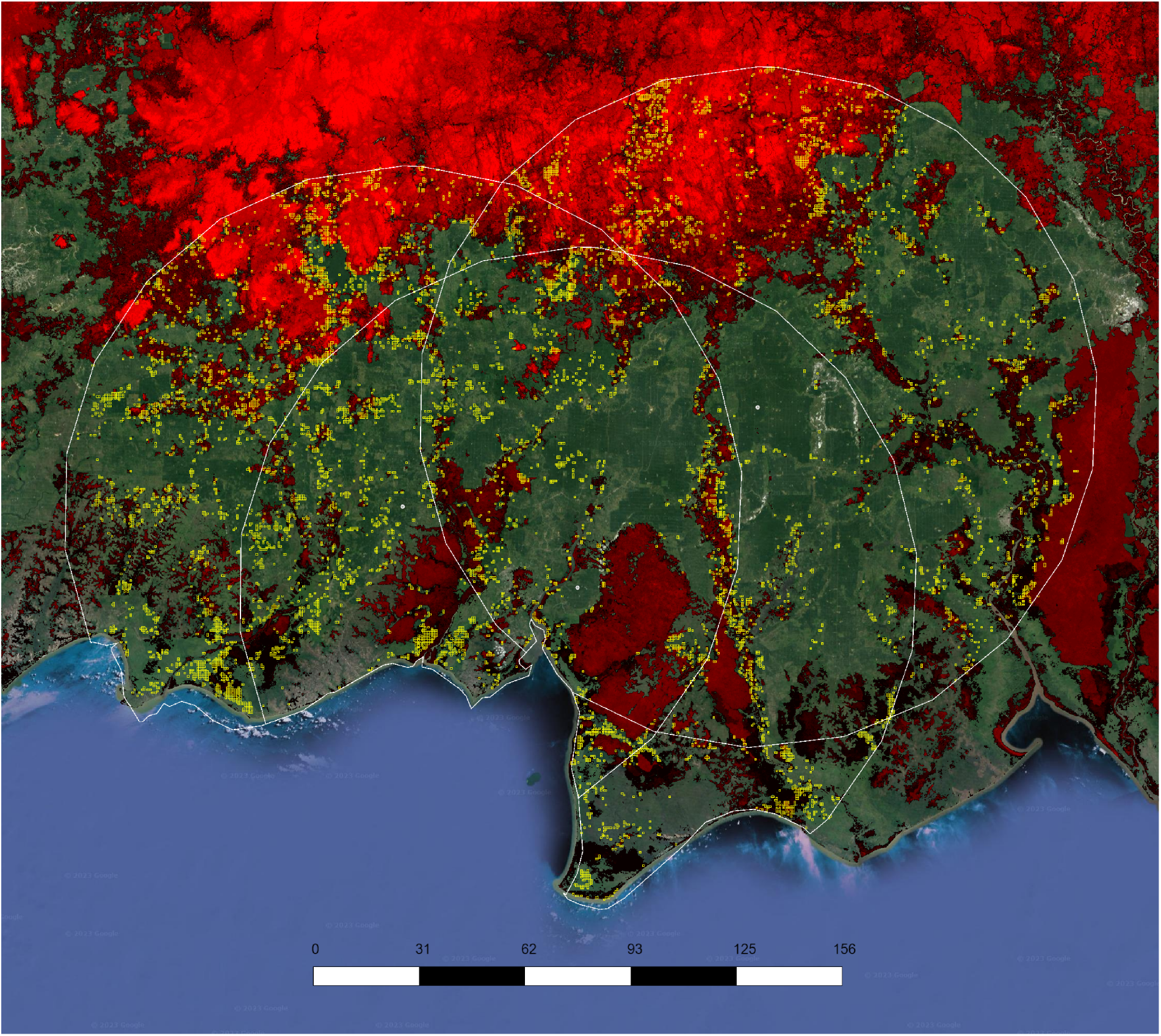
The spatial distribution of Forest Foresight hot zones (yellow) intersecting the prediction domains (white circular regions) of the three example mills (white markers) shown in Table 1. Per pixel carbon density is shown with a uniform color ramp from black (domain minimum = 0 MgC/pixel) to full red (domain maximum = 400 MgC/ha), over high resolution true color Google basemap imagery captured in year 2022 (© CNES, AirBus, Maxar). The green vegetative areas dominating all three prediction domains are predominantly industrial and small-holder palm plantations. Scale bar in kilometers.

## Discussion

By combining updated, high accuracy predictions of palm oil sourcing domains with machine learning-based deforestation predictions, we have demonstrated a method to proportionally allocate future deforestation, carbon loss, and biodiversity risk to relevant actors *ex ante*, thus enabling targeted mitigation activities before large-scale resource degradation has set in. In addition to risk mitigation within the *current* supply chain, these methods also make it possible for interested companies to continuously shape their supplier networks and volume origination portfolios to achieve longer-term goals, such as Scope 3 carbon emissions reductions in support of Net Zero commitments [27, 29, 30]. To enact large-scale change in agricultural value chains, this dynamic reshaping of physical commodity markets represents an important complement to the use of carbon credit schemes.

Recent research has offered diverse applications of machine learning and advanced modeling techniques to agricultural topics [79–82]. While this includes literature associated with palm oil supply chains [83–86], research has largely focused on making palm oil production more efficient [87]. With the exception of Joss [87], we are not aware of any published research that seeks to predict or evaluate the undisclosed sourcing domains utilized by palm oil suppliers. Joss relied on a cumulative cost sourcing model, as has been integrated here, exploring 8-, 24-, and 48-hour sourcing catchments based on a custom impedance dataset. Although the author was not able to test the reliability of her prediction domains, she too observed the gross lack of similarity between the industry standard radial approach and that informed by landscape impedance and real-world travel costs.

The species distribution modeling (SDM) framework we adopted has demonstrated moderate to high accuracy in other applications [62, 88], and our results indicate that the framework holds promise in our intended application. However, attempting to draw meaningful parallels between our findings and prior applications of SDM to non-human animal species, is questionable. A human’s capacity to alter its actions and environmental conditions based on higher order, rational thinking, leads to complex movement patterns that are only loosely analogous to the animal movement patterns typically driven by range establishment or maintenance, foraging, and reproductive behaviors. With the above in mind, the strength of our models can only be gauged with respect to our observed geolocational data. Where this data is itself derived from a traceability model [89], the real-world accuracy and utility of our predictions are inherently limited. Close inspection of our aggregated ping data reveals a consequential amount of data in areas that generally seem unfit for harvest, transportation, or aggregation of oil palm FFB (e.g., pings in residential neighborhoods). This raises obvious concerns, though we point to (a) the markedly improved accuracy of our models relative to the industry standard (albeit, also built from less than perfect training data), and (b) the distinction between palm origins (where palm FFB is harvested) and palm sourcing domains (where palm FFB are harvested, aggregated, transported, and processed).

Our intent in the present work was not solely to predict the locations of aggregated cellular pings of individuals harvesting palm FFB, but to predict the sourcing domains of Unilever’s in-network mills. A sourcing domain encompasses the locations where harvester pings are issued, as well as the intermediary space that supports transportation, aggregation, fallow fields, competitor exclusion, and expansion fronts, among others. In this way, a theoretical model trained solely on perfectly reliable real-world palm harvesting locations (palm origins), is not necessarily of greater utility, since such data would fail to capture the palm movement and larger activity space of palm oil suppliers. Future efforts would certainly benefit from stronger filtering workflows and an increase in data volume. To this end, we have begun piloting active traceability programs at select in-network mills, which we hope will provide an enhanced data model in the near future.

### Model considerations

Historically, palm FFB has commanded low cost per unit, and transportation over long distances has not, in our observation, been economically viable. However, the aggregate capacity of Indonesian palm processing facilities is twice as high as recent production volumes [87], potentially leading to high costs (e.g., [90]) and more distant sourcing than in the past. In the present work, we have chosen to make predictions at a maximum radial distance of 100 km (twice the industry standard). Changing this parameter would affect the ascription of risky behaviors by way of changing the allocation of responsibility at each pixelated location, though in most cases we do not believe the changes would be practically meaningful. The probability of sourcing for all mills is quite low at distances greater than 100 km, and for a given location, the rank order of suppliers with greatest responsibility is not apt to change. At the time of analysis, in Central Kalimantan, where our example mills are located, 46 groups own the 117 palm oil mills in Unilever’s previously disclosed supply chain. Of these, a subset of only nine own half of the mills, suggesting that large shifts in responsibility are unlikely when considering distances greater than 100 km.

### Industry adoption

As of year 2021, the palm oil industry was valued at USD $40 billion [91], and demand for palm oil is expected to increase [92]. Trends in palm-based deforestation showed a decline between 2017 and 2019, even reaching pre-2004 levels, but fluctuations in deforestation across time are correlated with the price of palm oil [90]. Where the current price has risen markedly over the past few years, in fact doubling since the start of the COVID-19 pandemic[90], there is now, as much as ever, a need for traceability in consumer goods supply chains. While certifying bodies, such as the Round Table on Sustainable Palm Oil (RSPO; [21, 22]) and the mandatory Indonesian Standard for Sustainable Palm Oil (ISPO; [23, 24] serve an important role in motivating audit-based traceability in the palm sector, they have been criticized for not addressing underlying causes of uncontrolled oil palm expansion [93, 94], in turn reflecting the need for additional traceability tools and technologies.

To this end, the results we have presented here show promise, though we still face a fundamental challenge of convincing those utilizing the current 50 km radial ascription scheme, to adopt something more complicated. The ubiquity of the current sourcing model, however unreliable, has provided participating suppliers and NGOs a certain level of familiarity and comfort. We have observed in the coordinated deployment of Forest Foresight, for example, that end users (primarily national governments) have demonstrated a keen interest in understanding the inner workings of even the most complex models, and that the explainability of new technologies is key in driving adoption. The broader adoption of more complicated data models will therefore require many NGOs and consumer goods companies to expend more time and resources on capacity building and stakeholder engagement than was previously thought necessary. Unilever and comparable consumer goods companies maintain active landscape- and jurisdictionally-oriented programs that are, by design, inclusive and managed by multi-stakeholder working groups (e.g., [95]). As traceability technologies continue to advance, these programs may play an increasingly important role in capacity building and adoption among key stakeholders.

In our analytical framework, we have proportionally ascribed predicted forest loss and associated carbon and biodiversity losses to the most-probable palm oil suppliers. However, these are not the only relevant factors driving forest loss and degradation. Our analyses do not directly illuminate post-conversion land use. Even in Indonesia and Malaysia, forest loss is not necessarily illegal or driven by growth in the palm oil industry, as predicted deforestation could be driven by legitimate logging interests, planned land use conversion in government delineated concessions, or natural fire, among others. In future adaptations of this work, it will be important to verify that the post-conversion land use of predicted (and realized) deforestation areas is actually tied to palm oil production.

## Methods

A complete and detailed description of our methods and justification for their use is presented in the Supplementary Information. All analysis was conducted with the Google Earth Engine (GEE) Python API [96]; Python data science and numerical computing libraries, including numpy [97], pandas [98, 99], sci-kit learn [100], and geemap [101], among others; the QGIS Python libraries [102]; and command line utilities to interface with the Google Cloud ecosystem, including the GEE Command Line Interface [103] and gsutil [104].

### Predicting sourcing domains

#### Training data

Our predictive models were based on observed geolocational data (hereafter, pings) obtained from Orbital Insights, who provided anonymized, aggregated data in the form of a marked point process. In our use-case, each ping represented the centroid of a 250 m x 250 m pixel, marked with the total number of device trips heading to and away from mills, the total number of unique device identifiers headed to and away from mills, coordinates, collection year, and facility to which each ping belongs (the parent mill). The unique facility identifiers were derived through data filtering workflows documented in Orbital Insight’s geolocation traceability model [89]. Our initial ping dataset was additionally filtered to include only those pings (a) in Indonesia, Malaysia, Brunei, and East Timor; (b) intersecting pixels containing *>*20% non-urban land cover (per [105]), (c) falling within mill prediction domains (see Predictive modeling framework), and (d) whose parent mill contained ≥ 50 aggregated pings, which we elected to use as a minimum threshold in modeling per [106]. After applying the above-mentioned filters, we had *n* = 622,015 observed locations (derived from 3,355,437 individual pings, 1,313,437 unique devices) to support models at 1,570 of 1,676 candidate mills.

### Predictive modeling framework

Predictive modeling was conducted on a mill-wise basis. For a given mill, the area over which predictions were made (the prediction domain), was determined by the closer of (a) 100 km (Euclidean) from the mill, (b) an international boundary, and (c) the coast of the land mass on which the mill fell. Within each mill-specific prediction domain, we implemented five different predictive models.

We approached analysis through the lens of species distribution modeling (SDM; [57, 58, 60, 107], where our “species of interest” was human palm workers. In this strategy, we augmented our presence-only data with background samples – also known as pseudo-absences [60, 108]. Allowing that a suitable number of background samples is a function of the landscape-level environmental conditions within which a species is found, we followed Phillips and Dudik [109] and Valavi *et al*. [62] in implementing a sensitivity analysis to optimize the sample size for our region and data. For each of 29 different background sample sizes between 25 and 50,000 locations (theoretical maximum is 502,524), we merged presence and background samples, fit an automatically tuned maximum entropy model (MaxEnt; [57, 58, 62, 106]) with default parameters, and then computed the area under the receiver operating characteristic curve (AUROC). This was repeated 10 times for each sample size, and AUROC was plotted against sample size (Fig. S1), leading us to select a sample size of 5,000 for all subsequent analyses (*n* = 8,471,035 labeled records across 1,570 mills).

### Explanatory variables

Traditionally, an SDM would exploit a potentially large collection of bioclimatic, edaphic, and topographic factors — the explanatory variables (predictors, covariates) — in modeling labeled data. Humans, however, represent an unusual species, in that we transcend, ignore, or otherwise modify these factors to go where we want and/or create what we want. In this way, common explanatory terms have limited bearing on our species’ distribution, or on the distribution of the palm worker subset. With this in mind, Table S1 presents our candidate set of raster-based explanatory variables intended to capture human use of the landscape. Using a subset of 21 mills in a tuning subset (see Accuracy assessment), we evaluated the utility of 16 combinations of predictors using 10-fold cross validation of AUROC derived from auto-tuned MaxEnt models (Fig. S4). In light of comparable model performance across candidate models (simplest model had an AUROC of over 91%), and increased visual interpretability, we elected to use four cost distance-based predictors in our models: cost distance; mean cost distance within 5 km radial neighborhoods; the difference in cost distance surfaces (cost delta) computed for the first and second closest mills to any given location; and the cost delta for first and third closest mills to any given location. Our expectation was that in locations where mills were closer together and exhibited greater resource competition, the cost delta values would be smaller, while in locations where the mills were more distant, cost deltas would be larger.

### Models

We explored four machine learning models: MaxEnt ([57–60], down-sampled random forest [61, 110], down-sampled gradient boosted regression trees (BRT) [63–65], and an ensemble of these three [66, 67]. These choices were predicated on Valavi et al’s [62] recent findings, which demonstrate that across a wide range of species (*n* = 225), geographies (*n* = 6 broad regions and many sub-regions), and sample sizes (*n* = 5-5,822), variations of these four models consistently out-performed many other parametric and non-parametric approaches. We also computed the industry standard model for each mill, permitting us to compare our predictions with the status quo.

MaxEnt does not suffer from imbalanced class distributions, and its application supported full use of our training data [57–60]. Random forest and BRT are both influenced by class imbalance, the effects of which were mitigated with down sampling. Here we followed Chen *et al*. [111], given the simplicity of their approach, its apparent performance characteristics [62, 110], and demonstrated success in other environmental modeling applications [112–114]. Weighting and sub-sampling are not possible using GEE’s default implementation of BRT (Haifeng 2014, Haifeng 2010), so our application is an approximation of the down-weighted BRT algorithm we were intending to emulate [62]; see Supplementary Information).

With respect to hyperparameter tuning, we relied on auto-tuned MaxEnt models, given Valavi et al’s [62] finding that manually tuned models showed no significant differences in performance (see also [109]). For our RF models, with the exception of the number of trees (B), we relied on default values supported by the literature ([62, 110, 115, 116]; bag fraction = 0.632, variables per split = √number of explanatory terms). Due to the bootstrapping and variable selection, the random forest algorithm is unlikely to overfit training data provided that *B* is sufficiently large for the out-of-bag (OOB) error to settle to a consistent rate [116–118]. To determine an appropriate number of trees to include in each forest, we derived the mean OOB error across *B* trees for 21 mills in a subsample of mills we refer to as our tuning subset (see Tuning subset, variable selection, and accuracy assessment below; Fig. S2; *B*=500). Unlike MaxEnt and random forests, BRT is particularly sensitive to hyperparameter tuning [119]. We sequentially tuned the number of trees (*n* trees = 1,000), the learning rate (shrinkage = 0.005), and the maximum number of leaf nodes (max nodes = 10), while using a sampling rate of 0.632 [110, 115](Fig. S3). All hyperparameter tuning was performed on our tuning subset.

We ensembled our individual models with an arithmetic mean [67]. In that all models predicted to the same [0-1] range of probability, the inputs were not rescaled. For our implementation of the industry standard model, we demarcated a 50 km (Euclidean) radial region around each mill, and classified any pings intersecting the region as class 1 (presences), and all others class 0 (absences). This model has no probabilistic equivalent that is currently employed, so comparisons with our probabilistic models were made in binary space based on confusion matrix derivatives.

### Tuning subset, accuracy assessment, and prediction

#### Tuning subset

Hyperparameter tuning and variable selection were based on a tuning subset of 21 of 1,570 mills (Table S2). These reflect a 0.5% stratified random sample of all mills, where strata were based on ping counts binned in increments of 250. The proportional representation of ping counts helped support generalizability in future extension of models to other mills.

#### Accuracy assessment

For each of our MaxEnt, random forest, gradient boosted regression tree, ensemble, and industry standard model workflows, we implemented 10-fold cross validation, training models on stratified random sample-based folds. The above-mentioned down-sampling routines were applied to these training folds. Trained models were applied to each test fold (*n* = 10) of each model (*n* = 5) for each mill (*n* = 1,570), producing a total of 78,500 sets of predictions. The accuracy and reliability of these predictions was assessed with AUROC, the true skill statistic (TSS), the AUC of precision-recall gain curves (AUPRG), precision-recall gain-based F_1_ accuracy (PRGF_1_), and Cohen’s kappa [76]. The supporting literature, context, and our rationale for selecting each of these metrics is described in the Supplementary Information. Our industry standard models were evaluated with TSS, PRGF_1_, and kappa only.

#### Statistical model comparison and prediction to grid

We were interested in a single modeling framework that, on average, yielded superior predictive results across all 1,570 mills. For this reason, we compared our models using one-way analysis of variance (ANOVA; [120]), which was applied independently to each of five sets of 1,570 model results associated with the above-mentioned accuracy assessment metrics. Where omnibus F-tests indicated significant differences in accuracy between model types, pairwise comparisons were performed with Tukey honest significant difference tests [121]. Our top performing model (ensemble) was then used to produce raster-based continuous valued predicted probability surfaces for each mill, extending across the mills-specific prediction domains.

### Predicting future deforestation events

Forest Foresight was launched by the World Wide Fund for Nature – Netherlands in Central Kalimantan, Indonesia, in 2018, as the “Early Warning System”, after development of an initial prototype in partnership with the Boston Consulting Group. The programme was expanded in 2019 to form a technical consortium with Deloitte, Amazon Web Services (AWS), Jheronimus Academy of Data Science (JADS) and Utrecht University, to 1) scale the predictive models to new landscapes; 2) improve the accuracy of the models; and 3) improve the usability of the tool. The predictive system has been piloted in Gabon, Suriname, and Indonesia (Kalimantan), while exploratory conversations are ongoing in Malaysia (Sarawak). A detailed technical description of the Forest Foresight program can be found in Van Stokkom *et al*. [55] as well as at https://forestforesight.atlassian.net/wiki/spaces/EWS/pages/33136/01+Introduction. We describe the main data and model features below.

### Labelled deforestation data

The primary data source for Forest Foresight is a set of labeled radar satellite images indicating deforestation event detections relative to a baseline. Throughout its operations, Forest Foresight has used two different sources of deforestation data, depending on regional availability: 1) the RADD alert data [122], and 2) deforestation alerts processed by SarVision [123], both of which are based on the freely available ESA Sentinel-1 data. The Sentinel-1 system currently consists of one operational satellite in a 12-day orbit, allowing for new deforestation detections on a bimonthly cadence. For Forest Foresight, deforestation alerts are obtained once a month at a 15-meter resolution. The data used in the current study were based on deforestation alerts provided by SarVision.

### Explanatory data and features

As predictive indicators for upcoming deforestation, labeled deforestation data is coupled with auxiliary geospatial datasets and spatial derivatives of the deforestation data itself (Table S1). Auxiliary predictors include distance to roads, waterways, coastlines [124], Euclidean distance to palm oil mills [125], gridded population density (WorldPop; [126]), slope, and elevation [127]. Spatial derivatives of the deforestation data include Euclidean distance to nearest recent deforestation patch younger than six months, edge density, patch density, aggregation, and landcover percentages within a one km^2^ focal neighborhood.

### Forest Foresight predictive modelling framework

These deforestation data and predictive features are combined in an XGBoost modelling framework [128] to predict areas at risk of deforestation in the next six months. Models are run at a resolution of 15 meters. A prediction threshold of ≥ 0.90 prediction probability is then applied to filter predictions of deforestation. Predictions above this threshold are retained and for reasons of generalizability and data sharing, subsequently down-sampled to 480 m “hot zones”, each indicating the proportional area of the hot zone above the prediction threshold, or at risk of deforestation. Resultant predictions at hot zone-level have a 70-80% user accuracy and a 40-50% detection rate. Additional details can be found in Van Stokkom *et al*. [55].

### Ascription of risk, carbon emissions, and biodiversity loss

We integrated our mill-level predicted sourcing surfaces with Forest Foresight predicted deforestation events using an array-based allocation approach. Our predicted sourcing surfaces overlapped one another for much of the study area, yielding a vector of probabilities at each coincident pixel. Working across surfaces, each pixel’s vector of probabilities was unit scaled to represent the relative share of risk ownership attributable to each mill. Future deforestation predicted by Forest Foresight was then allocated proportionally to each mill using the relative shares of ownership. This scheme was also used to ascribe carbon loss and biodiversity threat abatement measures to respective actors.

Our carbon data layer (30 m spatial resolution) was produced by Descrates Labs and included two core components: (1) A custom model of tree canopy height derived through random forest regression using explanatory terms extracted from NASA’s Global Ecosystem Dynamics Investigation Lidar (GEDI) collection [127], Sentinel-2 composites [129], latitude, and topographic derivatives ([130]; R2 = 0.62, MAE = 6.13m); and (2) ensembles of five regionally appropriate plot-level, tree height-dependent allometric equations for drylands and wetlands [131–133]. Uncertainty in tree canopy height predictions was propagated through the calculations of biomass to establish reliable non-parametric estimates of median carbon density per 30 m pixel. Ascription of carbon loss from deforestation events to relevant suppliers was based on Forest Foresight hot zones, using the following procedure:

1. Aggregate (sum) our 30 m carbon per pixel by hot zone.
2. For each hot zone, scale total carbon by the Forest Foresight estimated areal percent at risk.
3. Aggregate (mean) our 250 m resolution, unit scaled proportional allocations by hot zone.
4. Apportion the areally-adjusted carbon at risk by aggregated allocations.
5. Sum apportioned carbon loss by mill.

Risk to biodiversity was explored using the Integrated Biodiversity Assessment Tool’s (IBAT) Species Threat Abatement metric (STAR_*T*_; [78]). The STAR_*T*_ metric is designed to quantify the contribution of activities or operations at a given location, towards reduction in threat of species extinction [134]. The metric, which supports aggregation, cross-site comparison, and use at multiple scales, seeks to standardize and make measurable, nature-positive conservation action. We have elected to use START_*T*_ here given its broad applicability, its fundamental reliance on the IBAT database, and its direct support from the International Union for Conservation of Nature (IUCN). The STAR_*T*_ metric is effectively a weighted sum across all species within a given area of interest, where the proportion of a species’ population contained within the area of interest is weighted by the species’ IUCN Red List category (near threatened, vulnerable, endangered, and critically endangered) using established weighting ratios [78, 134, 135]. The STAR_*T*_ metric is presented as a reduction in threat of biodiversity loss, though it is effectively a biodiversity risk score, in that higher scores reflect areas with greater numbers of at-risk species, highly threatened species, or large portions of given species’ global population.

Ascription of STAR_*T*_ measures to relevant suppliers was based on Forest Foresight hot zones and mill-level sourcing areas using the following procedure:

1. Spatially disaggregate the global STAR_*T*_ dataset (4,961.46 m spatial resolution) by identifying the areal proportion that each hot zone is of the STAR_*T*_ pixel area on which it falls.
2. Scale the STAR_*T*_ value by each hot zone proportion, and also by the Forest Foresight estimated areal percent at risk.
3. Across coincident mills’ prediction surfaces, unit scale the mean predicted probability of sourcing within each hot zone, to obtain proportional allocations.
4. Apportion the areally-adjusted STAR_*T*_ at risk by proportional allocations.
5. Aggregate (sum) hot zone-level and mill-level STAR_*T*_ values (centiSTAR_*T*_ metrics).

## Supplementary information

A complete and detailed description of our methods and justification for their use can be found in the Supplementary Information.

## Acknowledgments

The authors extend their appreciation to Julia Chatterton, Ingrid Richardson, Martin Huxtable, Gullit Suwanto, and Jen Bennett for their comments and insights on the manuscript.

## Author contributions

A.W. and H.B.G. conceived the study; H.B.G., J.M.A, and J.T.-B. handled data curation; H.B.G., J.M.A., J.D., J.T.-B., and N.C. developed project methodology; H.B.G., J.T.-B., and M.V. performed formal analysis; H.B.G., J.M.A., J.D., J.T.-B., M.V., and A.W. were responsible for software development; H.B.G. and J.T.-B. validated the findings; J.D. and H.B.G. handled project administration; H.B.G., J.M.A., and J.T.-B. performed writing – original draft preparation; and all authors performed writing – review and editing.

## Software and data availability

The most accurate set of 1,570 predictions generated in this study are available under a CCBY-NC license. The GeoTiff image files, provided in WGS84 (EPSG:4326) with a nominal (equatorial) spatial resolution of 250 m, can be accessed through Zenodo at https://doi.org/10.5281/zenodo.7766561. The corresponding images are also currently available as a Google Earth Engine asset in the coordinate reference system used during production, which was a World Mollweide projection (ESRI:54009) with a modified central meridian (longitude of origin) of 109.5 degrees, and a spatial resolution of 250 m. The Google Earth Engine ImageCollection is available at https://code.earthengine.google.com/?asset=projects/ul-gs-d-901791-09-prj/assets/users/hglick/Glick_et_al_2023/Ensemble.

