## Supplementary Information for "Model-based prediction and ascription of deforestation risk within commodity sourcing domains: Improving traceability in the palm oil supply chain"

[1]NGIS Australia, 1a/53 Burswood Road, Burswood, WA, 6100, Australia.
[2]Unilever, 800 Sylvan Avenue, Englewood Cliffs, 07632, NJ, USA.
[3]World Wide Fund for Nature Netherlands, Driebergseweg 10, 3708 JB, Zeist, The Netherlands.
[4]The Nature Conservancy, 4245 North Fairfax Drive, Suite 100, Arlington, VA, 22203.
[5]Sustainable Sourcing, Unilever, Unilever Campus, 18 Nepal Park, Singapore, 139407.
[6]Google LLC, 1600 Amphitheatre Parkway, Mountain View, 94043, CA, USA.

Contributing authors:;

**Abstract**

Palm oil accounts for approximately 50% of global vegetable oil production, and trends in consumption have driven large-scale expansion of oil palm (*Elaeis guineensis*) plantations in Southeast Asia. This expansion has led to deforestation and other socio-environmental concerns that challenge consumer goods companies to meet no deforestation and sustainability commitments. In support of these commitments and supply chain traceability, we seek to improve on the current industry standard sourcing model for ascribing social and environmental risks to particular actors. Using passive geolocational traceability data ($\boldsymbol{n}$ = 3,355,437 cellular pings) and machine learning models, we evaluate the industry standard sourcing model, and we predict with high accuracy the undisclosed sourcing domains for 1,570 Indonesian and Malaysian palm oil mills on the Universal Mill List (as of November 2021). In combination with the World Wide Fund for Nature – Netherlands' Forest Foresight, we use our predicted sourcing domains to provide an illustrative example of the proportional allocation of future deforestation, carbon loss, and biodiversity risk to relevant actors, permitting targeted outreach, contract negotiation, and mitigation of large-scale resource degradation. This example depends on a subset of model predictions in the absence of disclosed traceability data. The utilization of additional predictions or disclosed traceability data would influence and improve the results.

**Keywords:** *Elaeis guineensis*, palm oil, risk assessment, deforestation, NDPE, supply chain, machine learning

### Methods

In support of clarity, we detail our methods in the following subsections:

1 Predicting sourcing domains

1.1 Training data
1.2 Predictive modeling framework
1.3 Explanatory variables
1.4 Models

1.4.1 MaxEnt
1.4.2 Down-sampled random forests
1.4.3 Down-sampled gradient boosted regression trees
1.4.4 Ensemble
1.4.5 Industry standard

1.5 Tuning subset, variable selection, and accuracy assessment

1.5.1 Tuning subset
1.5.2 Variable selection
1.5.3 Accuracy assessment

1.5.3.1 AUROC and TSS
1.5.3.2 AUPRG and $F_1$
1.5.3.3 Kappa
1.5.3.4 Statistical comparison and prediction to grid

2 Ascription of risk, carbon emissions, and biodiversity loss

All analysis was conducted with the Google Earth Engine (GEE) Python API [1](earthengine-api v. 0.1.316); Python data science and numerical computing libraries, including numpy [2](v. 1.22.1), pandas [3, 4](v. 1.4.0), sci-kit learn [5](v. 1.4.0), and geemap [6](0.14.0), among others; the QGIS Python libraries [7](v. 3.16); and command line utilities to interface with the Google Cloud ecosystem, including the GEE Command Line Interface [8](within earthengine-api v. 0.1.316) and gsutil [9](v. 5.10).

### 1 Predicting sourcing domains

The area from which a palm processing facility (hereafter, palm mill) sources raw FFB, has generally been estimated using the 50 km radial sourcing model described in the main text. This industry standard means of ascription fails to adapt to variable topographic or transportation limitations, and fails to address fundamental issues brought by overlapping sourcing domains. It neither draws on the available sourcing context (e.g., palm plantations, human settlement patterns) nor offers any measure of reliability. To our knowledge, this approach, however ubiquitous, has never been properly evaluated, and ultimately provides a false sense of understanding. Here we put forth a strategy for improving upon the radial sourcing model, drawing on empirical geolocational traceability data and the application of non-parametric machine learning methods to generate spatially explicit rasterized probability surfaces, where each pixel reflects the probability that it falls within a given mill's sourcing domain.

#### 1.1 Training data

Our predictive models were based on observed geolocational data (hereafter, pings) obtained from Orbital Insights. These locations represent anonymized and aggregated passive geolocation data from any individual who (a) owns a cellular device (i.e., cellular telephone, tablet), (b) whose device has one or more applications that record geolocational data using a built-in GPS transceiver (many applications), and (c) who has not deliberately turned off geolocational awareness on all of their applications. When a cellular device makes contact with cellular and/or satellite reference stations, it transmits a packet of information to the service provider to document its location. Depending on the topography, number of cellular towers and/or satellites, and quality of the device transceiver, the transmitted coordinates will be more or less locationally accurate. Older cellular devices generally produce less accurate locational data. Orbital Insights collects ping data from across the globe, and estimates a 2% penetration rate across Southeast Asia (personal communication, A. Douglas Sept. 2021). This means that Orbital Insights acquires and processes data from only 2% of cellular device owners in the region, either because the remaining population has geolocational services turned off, or because there is another limiting factor (e.g., landscape, device age, physical remoteness).

Orbital Insights provided anonymized, aggregated geolocation data in the form of a marked point process. In our use-case, each geolocated ping represented the centroid of a 250 m x 250 m pixel, marked with the total number of device trips heading to and away from mills, the total number of unique device identifiers headed to and away from mills, coordinates, collection year, and facility to which each ping belonged (the parent mill). The unique facility identifiers were derived through data filtering workflows documented in Orbital Insight's geolocation traceability model [10]. Among other filters, the raw, non-aggregated pings are isolated or tagged based on locational accuracy, travel speed, residence time, and visitation to facilities of interest. Aggregated pings represent a collection of devices whose observed behavior ties them to one or more specific facilities, and which supports their use as model training data.

Our initial ping dataset contained $n$ = 1,484,899 aggregated locations (years 2019 and 2020), comprised of 5,516,924 individual pings across 2,736,614 unique devices. We subsequently filtered the raw data to include only those (a) in Indonesia, Malaysia, Brunei, and East Timor; (b) intersecting pixels containing >20% non-urban land cover (per [11]), (c) falling within mill prediction domains (see §Predictive modeling framework), and (d) whose parent mill contained ≥50 aggregated pings, which we elected to use as a minimum threshold in modeling. Small sample sizes are common in species distribution modeling (SDM), particularly when working with rare species. Wisz *et al.* ([12]) evaluated several models across typical record counts ($n$ = 10, 30, 100) and found that while some models performed reasonably with small samples, none of the tested algorithms performed consistently well with <30 records. After applying the abovementioned filters, we had $n$ = 622,015 observed locations (3,355,437 individual pings, 1,313,437 unique devices) to support models at 1,570 of 1,676 mills in Unilever's supply chain.

#### 1.2 Predictive modeling framework

Predictive modeling was conducted on a mill-wise basis. For a given mill, the area over which predictions were made (the prediction domain), was determined by the closer of (a) 100 km (Euclidean) from the mill, (b) an international boundary, and (c) the coast of the land mass on which the mill fell. Palm is subject to high levies and strict export control in Southeast Asia, and the value of palm FFB is generally too low to warrant transportation across major bodies of water. The 100 km represents a conservative transportation distance (2 × industry standard). Within each mill-specific prediction domain, we implemented six different models using two broad strategies to convert our presence-only data into binary data suitable for supervised classification and regression tasks. We detail one strategy and five models here, and offer a brief summary of a second strategy and sixth model used in pilot testing.

We approached analysis through the lens of species distribution modeling (SDM; [13–16]), where our species of interest was human palm workers. In this strategy, we augmented our presence-only data with background samples – also known as pseudo-absences [16, 17]. The term "background samples" reflects the intention: a spatially random sample of the background environmental conditions within which your species lives, irrespective of the locations of the observed data [18, 19]. The number of background samples used in published SDMs is not fixed. An influential benchmark study used 10,000 locations [20], though some research indicates that a larger number of samples is required for optimal model performance and convergence with presence-only point process models [21]. Allowing that a suitable number of background samples is a function of the landscape-level environmental conditions within which a species is found, we followed Phillips and Dudik [22] and Valavi *et al.* [19] in implementing a sensitivity analysis to optimize the sample size for our region and data. Here, the logic is that as the background sample becomes sufficiently large, the variability inherent in the environmental conditions is suitably represented, and separability between classes is maximally stable for the data at hand [19]. We conducted this analysis using a maximum entropy model [MaxEnt; 13, 14] because (a) it was one of the models we had chosen for prediction, (b) it is not a stochastic model that leads to highly variable results [13, 14], (c) it performs more consistently than many other models with small sample sizes [12], and (d) it is computationally fast to implement [19].

We selected two mills that fell entirely within their parent landmasses (i.e., having maximal prediction domains), which fell in common sourcing territory, and which each had a moderate number of associated pings ($n = \approx 500$). For each of 29 different background sample sizes between 25 and 50,000 locations (theoretical maximum is 502,524), we merged presence and background samples, fit an automatically tuned MaxEnt model with default parameters, and then computed the area under the receiver operating characteristic curve (AUROC). This was repeated 10 times for each sample size. Fig. S1 details the asymptotic results for one mill (second omitted out of similarity), which indicates diminishing returns after a sample of size about 1,000. We elected to use a sample of size 5,000 for all subsequent analyses, striking a conservative balance between computation time and reduction in variability. Our final collection of presence-absence locations had $n = 8{,}471{,}035$ labeled records across 1,570 mills.

##### 1.3 Explanatory variables

Traditionally, an SDM would exploit a potentially large collection of bioclimatic, edaphic, and topographic factors — the explanatory variables (predictors, covariates) — in modeling labeled data. Humans, however, represent an unusual species, in that we transcend, ignore, or otherwise modify these factors to go where we want and/or create what we want. In this way, common explanatory terms, such as soil quality, precipitation regimes, insolation, or terrain ruggedness have limited bearing on our species' distribution, or on the distribution of the palm worker subset. With this in mind, Table S1 presents our candidate set of raster-based explanatory variables intended to capture human use of the landscape. Two require further explanation.

The two cost delta surfaces were included to capture resource competition between mills. In practice, a cost distance surface was generated for each mill, and the unique mill identifier was attached to each pixel as a two-element array. We then assembled the full collection of cost distance array surfaces, treating them as a large multidimensional data cube, and ordered spatially intersecting elements. This operation individually ordered each pixel by the collection of cost distance values present at that pixel, carrying along the respective identifiers during the ordering. In the resulting data product, the first element at each location was the cost distance to the nearest mill, along with that mill's identifier. The second element was the cost distance to the second nearest mill, along with its identifier, followed in turn by the third, fourth, etc. We used this product to compute the difference between the travel costs to the first and second closest mills, and between the first and third closest mills. Here, our expectation was that in locations where mills were closer together and exhibited greater resource competition, the cost delta values would be smaller, while in locations where the mills were more distant, cost deltas would be larger.

For each of the 1,570 mills, the covariates detailed in Table S1 were created, numerically standardized, and and clipped to the mill-specific prediction domains using a spatial resolution of 250 m and a customized World Mollweide projection (ESRI:54009) in which the central meridian was shifted to 109.5 degrees (approximate center of our study area). Aggregation of higher resolution data products used appropriate mean, modal, or summation reducers, while disaggregation employed direct up-sampling, dividing larger pixels into like-valued smaller pixels. We then used standard extraction tools to bind covariate values to the 8,471,035 presence-absence locations.

##### 1.4 Models

We explored four machine learning models: MaxEnt, down-sampled random forest, down-sample gradient boosted regression trees, and an ensemble of these three. These choices were predicated on Valavi et al's [19] recent findings, which demonstrate that across a wide range of species ($n = 225$), geographies ($n = 6$ broad regions and many sub-regions), and sample sizes ($n$ = 5-5,822), variations of these four models consistently outperformed many others, including both common parametric (e.g., generalized linear models, ridge regression) and non-parametric machine learning models (e.g., unbiased conditional inference forests, XGBoost, support vector machines). The apparent predictive strength paired with strong generalizability made these appropriate choices for our application. We also computed the industry standard model for each mill, permitting us to compare our predictions with the status quo.

**Table S1**: Candidate set of raster-based explanatory variables explored in predictive modeling.

| Variable | Intended to capture | Depends on | Scale (m) | Notes |
|---|---|---|---|---|
| Cost distance | Infrastructural and temporal limitations to movement | [23] | 927.67 | Cumulative cost applied to [23] in GEE catalog. |
| Mean cost distance | Generlized version of above | Cost distance, above | 250 | 5 km radial focal mean of above. Intended to smooth potential errors in [23]. |
| Palm mask | Raw commodity of interest | Proprietary | 20 | CNN trained on 8,000 labeled locations using Sentinel-1 GRD, surface reflectance composites, and SRTM terrain data. Discretized with 0.5 decision threshold. F1 accuracy = 0.91 |
| Distance to palm | Logistical limitations in harvesting and transportation | Palm mask, above | 250 | Euclidean distance to palm |
| Urban-rural mask | Human settlement and activity | [11, 24] | 100 | Copernicus Global Land Service (CGLS) Dynamic Landcover Map (CGLS-LC100). Isolation of “Settlement Area”, “Airport/Harbour”, and “Transmigration Area from [24]; and “Urban/built up” from [11]. Land cover circa 2006, reclassified from Indonesian Ministry of Forestry (2009; 1:250,000). |
| Distance to urban centers | Logistics related to human settlement and activity | urban-rural mask, above | 250 | Euclidean distance to isolated urban portion of urban-rural mask, above. |
| Population density | Population density | [25] | 30.92 | Fusion of machine learning-based image analysis and census data. Delivered as people per pixel. |
| Road density (all) | Transportation infrastructure | [26, 27] | 250 | 10 km radial focal mean of rasterized fusion of (a) all OSM roads as of 20220107, and (b) Facebook’s AI-assisted road tracings, which were rasterized at 10 m. |
| Road density (tertiary +) | Potentially more realistic transportation infrastructure | [26, 27] | 250 | 10 km radial focal mean of rasterized fusion of (a) OSM roads size tertiary or larger, as of 20220107, and (b) Facebook’s AI-assisted road tracings, which were rasterized at 10 m. |
| Cost delta 1-2 | Competition between suppliers | [23] | 250 | Difference in cost distance values between first and second closest mills, for all locations. |
| Cost delta 1-3 | Competition between suppliers | [23] | 250 | Difference in cost distance values between first and third closest mills, for all locations. |

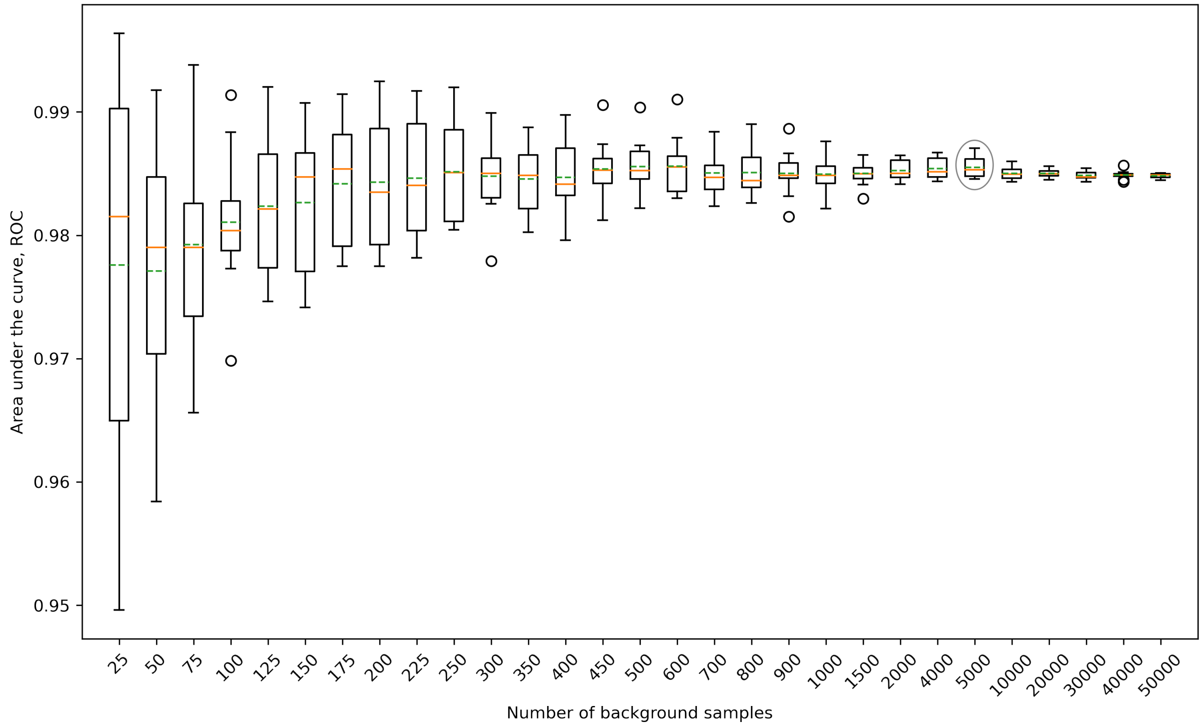


**Fig. S1**: The number of random background samples (pseudo-absences) and the associated area under receiver operating characteristic curve (AUROC) for auto-tuned MaxEnt models applied to a representative mill. Each box represents 10 iterations with unique random background samples. As the number of background samples increases, we observe an asymptotic increase in AUROC, and a concomitant reduction in variability thereof. Theoretical maximum sample size is 502,524 samples. Boxes cover the first through third quantiles (hinges), whiskers of 1.5×interquartile range, and further outliers (points). Median and mean values are shown in green and orange, respectively.

##### 1.4.1 MaxEnt

There are well over 1,000 published applications of the MaxEnt model to species distribution and environmental niche modeling since 2006 [16], and the mechanics of the model have been detailed elsewhere [13, 14, 16, 28]. In practice, we employed the GEE `ee.Classifier.amnhMaxEnt` function (probability output mode), which is a wrapper for the open-source Java implementation of the algorithm in the Statistical and Machine Intelligence Learning Engine (SMILE) library (https://github.com/haifengl/smile) [29, 30]. We elected to use auto-tuning functionality built into this algorithm, so as to avoid feature engineering and tuning of regularization hyperparameters. This choice was based on Valavi et al's [19] findings, which indicated that across many similar applications, there was no statistically significant difference between manually tuned MaxEnt models and those relying on the default parameters. This finding may be a function of the original data used to create the default parameters [22], which was essentially the same as that used by Valavi *et al.* [19]. However, even in the original model parameterization process, Phillips and Dudik [22] found that their default tunings performed nearly as well as models tuned on evaluation data. Avoiding model tuning was desirable, in that the GEE architecture and default object structures [1] are not as conducive to certain types of cross validation and grid search strategies as are other numerical computing frameworks. In addition, MaxEnt's predictive strength does not suffer from imbalanced class distributions, as do other models that benefit from cost sensitive learning (e.g., [31–33]). We made full use of the presence-absence data, without down-sampling or other cost sensitivity adjustments.

##### 1.4.2 Down-sampled random forest

Class imbalance in training data is a persistent challenge in many real-world settings [34–37]. A number of approaches have been used to improve classifier performance and consistency when applied to imbalanced data, including cost sensitive learning [31–33], various sampling schemes [34, 38–40], and ensemble learning methods [34, 36], among others. As an ensemble learner, the ubiquitous random forest algorithm [19, 41] may generalize better [42] and be less sensitive to class imbalance ([36, 43], c.f. [44]) than the classification and regression trees (CART, [45]) on which the algorithm depends. But where the component decision trees are known to be sensitive to class imbalance [34], a number of random forest variations have been developed to mitigate the potential ill effects [46]. Here we have elected to pair random forests with a down-sampling approach proposed by Chen *et al.* [38], given its simplicity, apparent performance characteristics [19, 44], and demonstrated success in other environmental modeling applications [47–49].

In down-sampled random forest, or balanced random forest [38], the analyst iteratively balances the majority and minority classes by sub-sampling the majority class. If balancing is performed only once, prior to running the random forest algorithm, there might be loss of information from the ensemble as a whole, given that a potentially large portion of the majority class would be entirely omitted [38]. To circumvent this issue, a bootstrapped sample (with replacement) is taken of both classes for each tree in the forest, creating an ensemble of many different balanced trees. In practice, we used Google Earth Engine's `ee.Classifier.smileRandomForest` function (probability output mode), which is a wrapper for the random forest algorithm available in the SMILE library [29, 50]. Weighting and sub-sampling are not possible using the default implementation of the algorithm, so for each mill we produced $B$ down-sampled single-tree "forests", used the resulting $B$ trees to predict to raster grid format, and then computed pixel-wise means across the coincident $B$ grids.

To avoid extensive hyperparameter tuning in GEE, we used common software default values shown to perform well in many circumstances. We permitted an unbounded number of leaf nodes in each tree, each with a minimum required population of one element, potentially leading to deep trees; we used a bag fraction of 0.632, which is the empirical proportion of original data points supporting bootstrap samples [44, 51]; and we used $\sqrt{\text{number of explanatory variables}}$ as the number of variables per split [19, 52]. Due to the bootstrapping and variable selection, random forests is unlikely to overfit training data provided that $B$ is sufficiently large for the out-of-bag (OOB) error to settle to a consistent rate [52–54]. To determine an appropriate number of trees to include in each forest, we derived the mean OOB error across $B$ trees for 21 mills in a subsample of mills we refer to as our tuning subset (see §Tuning subset, variable selection, and accuracy assessment below). Figure S2 shows relative consistency in the OOB error and the variability of that error across a wide range of tree sizes. We selected a $B$ of 100 trees.

##### 1.4.3 Gradient boosted regression trees

The gradient boosted regression tree algorithm [55–57] (hereafter BRT) is not dissimilar from random forests in its use of a large number of classification and regression trees. However, where random forests ensembles a large collection of decorrelated and potentially deep trees built on bootstrapped samples, boosted trees are shallow trees built sequentially on iteratively modified versions of the original input data [52]. Conceptually, the BRT algorithm relies on Valiant's "probably approximately correct" framework [58, 59], in which there is high probability that the model will have low generalization error, obtained through the use of many weaker learners. The BRT algorithm implements a stage-wise modeling approach, fitting each sequential CART to the previous stage's residuals [59, 60]. This general approach endeavors to "boost" the previous stage's results by iteratively reducing the gradient of the selected loss function [59, 60]. Boosted regression trees have been uncommon in ecology [60], and by extension, SDMs.

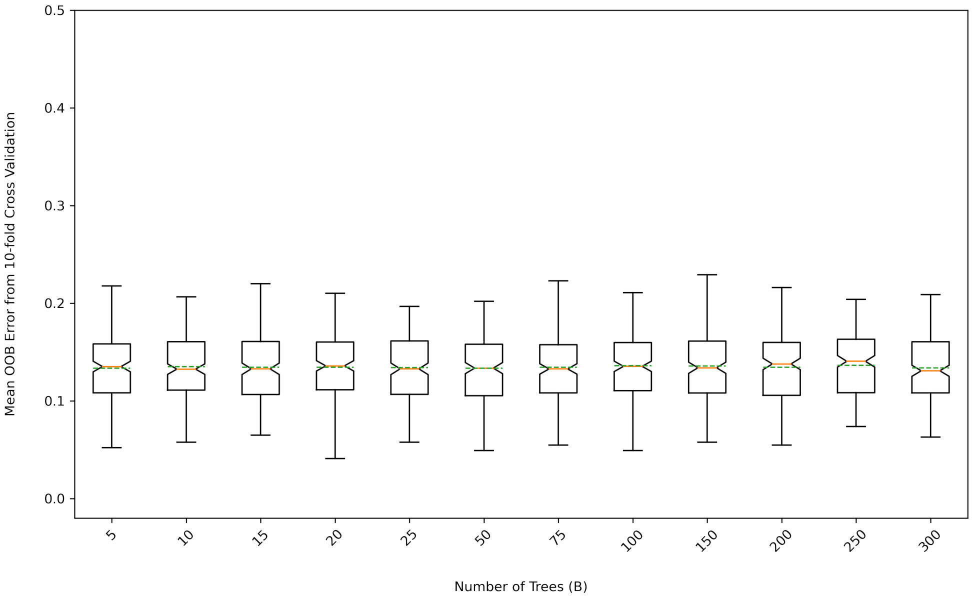


**Fig. S2**: 10-fold cross validated out of bag error for down-sampled random forests models (all candidate variables), by increasing tree count (B). Each box represents mean out of bag error from 10-fold cross validation for 21 mills in a tuning subset. Boxes cover the first through third quantiles (hinges), whiskers of 1.5*interquartile range, and further outliers (points). Median and mean values are shown in green and orange, respectively.

In practice, we used Google Earth Engine's `ee.Classifier.smileGradientTreeBoost` function (probability output mode), which is a wrapper for the GradientTreeBoost algorithm available in the SMILE library [29, 61]. Following Valavi *et al.* [19], we intended to implement a down-weighted BRT algorithm to accommodate gross class imbalance. In down-weighted BRT, records from the majority class are iteratively down-weighted to collectively carry equivalent influence as the minority class [20]. This is not possible in GEE, so we elected to use down-sampling [38] as we had in random forest modeling. In the case of random forests, down-sampling can occur once for each tree. In the case of BRT, the stage-wise evolution does not lend itself to this approach, since all shallow trees for a given model are built from the same input data. For this reason, our 10-fold cross validated results (see §Tuning subset, variable selection, and accuracy assessment) reflect only an approximation of down-weighted BRT, since each of the 10 models is effectively a balanced model in which we limited the number of original background samples.

Unlike MaxEnt and random forests, BRT is particularly sensitive to hyperparameter tuning [60]. Using Python's sci-kit learn library [5], we sequentially tuned the number of trees (n.trees), the learning rate (shrinkage; start value = 0.005), and the maximum number of leaf nodes (max.nodes; start value = 8). These hyperparameters are, with the exception of sampling rate (fixed value = 0.632, see §Down-sampled random forests), the only parameters exposed in GEE's utility. The minimum samples per split was held constant at 6 in support of generality and shallow tree growth, though this parameter is not controllable in GEE. Each tuning process was conducted on the 21 mills in our tuning subset (see §Tuning subset, variable selection, and accuracy assessment), and was subjected to 10-fold cross validation. Where some loss functions are more sensitive to noisy data than are others, we used least absolute deviation in both tuning and application, which is more robust to noise than traditional least squares [42, 60]. We acknowledge that different implementations of the algorithm (Python vs Java) may result in slightly different findings.

From Fig. S3a we selected n.tree = 1000 for all subsequent analyses. Figure S3b reflects a grid search over shrinkage values of [0.001, 0.0025, 0.005, 0.0075, 0.01, 0.025, 0.05, 0.075, 0.1]. From these results we selected a suitable learning rate of 0.005 for all subsequent analyses. Friedman [56] reflects on how decreasing the shrinkage value increases the optimal value for n.trees, so this rate is particular to our choice of n.trees = 1000. Figure S3c reflects tuning of max.nodes, which is, to some extent, a measure of tree depth. For this reason, tree depth is usually constrained to a small value, and in GEE the default is max.nodes = 6. If we permit too many max.nodes, we risk limiting generalization. From Fig. S3c we selected max.nodes = 10, which is, in the case of a symmetrical tree, somewhere between three and four levels deep (i.e., still shallow).

##### 1.4.4 Ensemble

Ensemble modeling, or consensus modeling, is the process of combining the models or results from two or more individual models, and is predicated on the idea that the combination of individual models will yield stronger results than any individual model. This is not dissimilar to contemporary ensemble learners appearing in the machine learning literature. Ensembling approaches and recognition of its benefits have existed for decades [62–66]. Ensembling is common in SDM [67], where it has been shown to yield greater and more robust accuracies than individual models [19, 66, 68]. Valavi *et al.* [19] found that an ensemble of their top performing models, including rescaled and averaged Generalized Additive Models, lasso regressions, MaxEnts, down-weighted BRTs, and down-sampled random forests, performed better than any individual model. We follow this approach here, and for each mill we computed an arithmetic mean [68] of predictions from our MaxEnt, down-sampled random forest, and down-sampled BRT models. In that all models predicted to the same [0-1] range of probability, inputs were not rescaled.

##### 1.4.5 Industry standard

For each mill, we produced and evaluated the industry standard 50 km radial sourcing model. This was achieved by demarcating a 50 km (Euclidean) radial region around each mill, and classifying each of a mill's pings intersecting that circular region as class 1 (belonging to the mill), and each of the points >50 km from the mill as class 0 (not belonging to the mill). This model has no probabilistic equivalent that is currently employed, so comparisons with our probabilistic models were made in binary space based on confusion matrix derivatives.

#### 1.5 Tuning subset, variable selection, and accuracy assessment

##### 1.5.1 Tuning subset

Hyperparameter tuning and variable selection were based on a tuning subset of 21 of 1,570 mills (Table S2). These reflect a 0.5% stratified random sample of all mills, where strata were based on ping counts binned in increments of 250. The proportional representation of ping counts helped support generalizability when extending models to other mills.

**Table S2**: Our tuning subset of 21 mills used in variable selection and model tuning workflows. This selection was derived through stratified random sampling of all mills, where strata were based on aggregated cellular ping counts.

| Mill | Count | Stratum | Mill | Count | Stratum |
|---|---|---|---|---|---|
| 1 | 5216 | 5000 | 12 | 6556 | 6500 |
| 2 | 5180 | 5000 | 13 | 6821 | 6750 |
| 1 | 5216 | 5000 | 12 | 6556 | 6500 |
| 2 | 5180 | 5000 | 13 | 6821 | 6750 |
| 3 | 5184 | 5000 | 14 | 7059 | 7000 |
| 4 | 5165 | 5000 | 15 | 7356 | 7250 |
| 5 | 5274 | 5250 | 16 | 7513 | 7500 |
| 6 | 5252 | 5250 | 17 | 7939 | 7750 |
| 7 | 5276 | 5250 | 18 | 8046 | 8000 |
| 8 | 5656 | 5500 | 19 | 9492 | 9250 |
| 9 | 5902 | 5750 | 20 | 9503 | 9500 |
| 10 | 6089 | 6000 | 21 | 11042 | 11000 |
| 11 | 6285 | 6250 | | | |

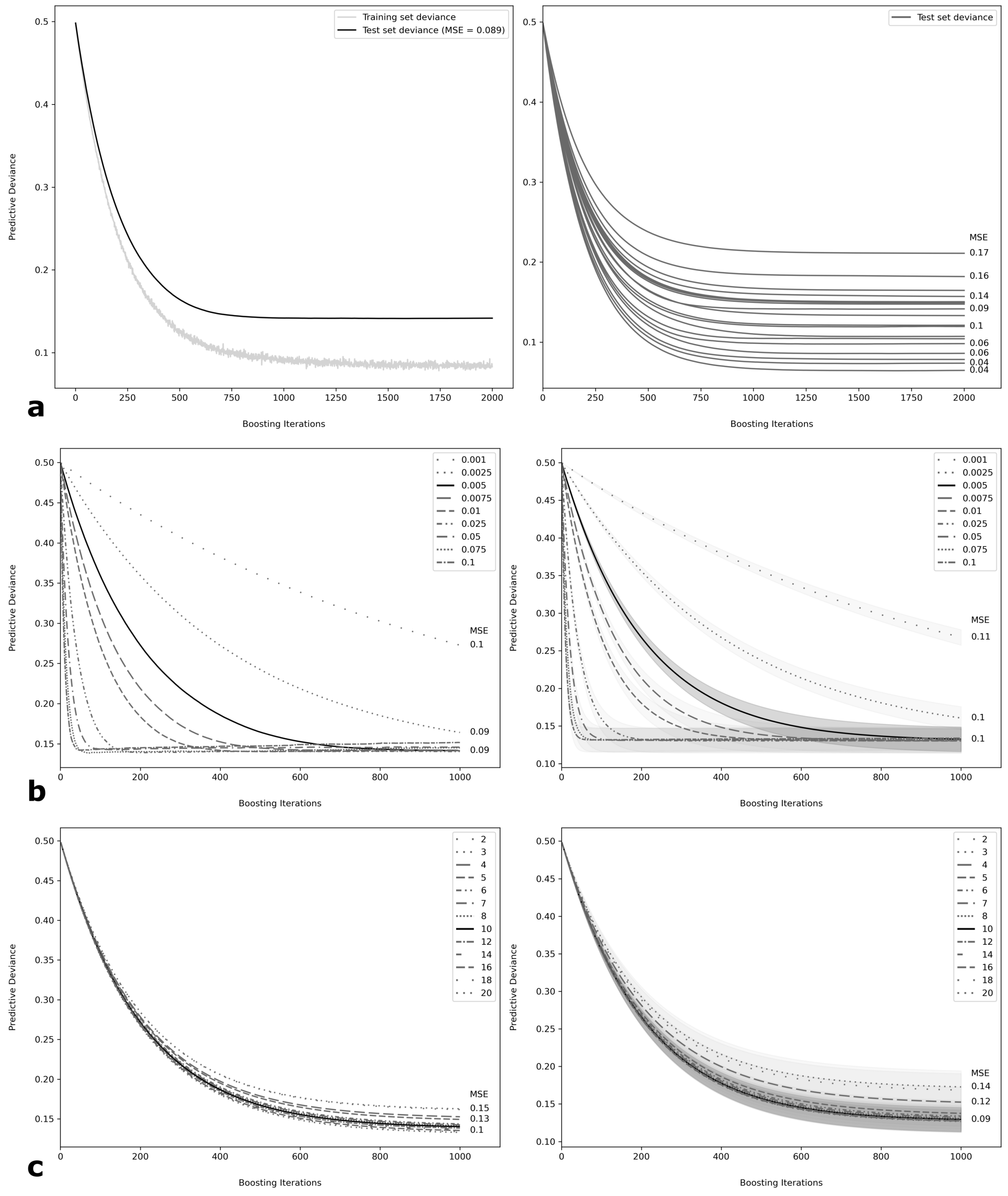


**Fig. S3**: Results of hyperparameter tuning of down-sampled gradient boosted regression tree models applied to all candidate variables. Row a = tuning for number of trees (n.tree); row b= tuning for learning rate (shrinkage); row c = tuning for maximum number of tree nodes (max.nodes). Plots in the left column provide examples of tuning results for a single mill. Plots in the right column contain the means (dark lines) of 10-fold cross validated model mean values for each of 21 mills in our tuning subset, along with their 95% confidence intervals (shaded areas). Except where noted, all lines are built from test sets. MSE = mean squared error.

##### 1.5.2 Variable selection

We selectively evaluated variable importance across our tuning subset, ultimately electing to use a limited number of our explanatory terms. Based on preliminary testing and inherent interest, we retained in each model the four cost distance-based terms (cost distance, mean cost distance, cost delta 1-2, cost delta 1-3). To these we added the 15 combinations of remaining explanatory terms, and computed 10-fold cross validated AUROC from auto-tuned MaxEnt models applied to each of the 21 mills in the tuning subset (Fig. S4). While Fig. S4 shows statistically significant differences between models when looking across the tuning subset, even the most restrictive model (model A) had a mean AUROC of over 91%. Model B, which included a single additional predictor, had superior performance, but the predicted domains for B and other more complex models, had enough visual discontinuity that we questioned downstream industry adoption. For this reason, we relied on variable subset A for all subsequent modeling.

##### 1.5.3 Accuracy assessment

For each of our MaxEnt, random forest, gradient boosted regression tree, and ensemble workflows, we implemented 10-fold cross validation, training models on stratified random sample-based folds. The above-mentioned down-sampling routines were applied to these training folds. Trained models were applied to each test fold ($n$ = 10) of each model ($n$ = 4) for each mill ($n$ =1,570), producing a total of 62,800 sets of predictions. The accuracy and reliability of these predictions was assessed with AUROC, the true skill statistic (TSS), the AUC of precision-recall gain curves (AUPRG), recision-recall gain-based $F_1$ accuracy, and Cohen's kappa. While the context and our rationale for selecting each of these metrics is briefly described below, we acknowledge that there is an almost bewildering number of published evaluation metrics. In the context of SDM, the literature is contradictory, indicating both support and criticism for most of the common metrics. In recognition that SDM is still dominated by metrics that have been repeatedly criticized [69], we have selected some that are familiar, maximize information retention, permit model comparison, and which receive support from various camps.

###### *1.5.3.1 AUROC and TSS*

There is growing evidence that in the context of SDM, threshold-based accuracy metrics are overly restrictive and diminish the information content in the final predictions [19, 70]. For this reason, we utilized two threshold-independent curve-based metics. Receiver operating characteristic curves are a common model evaluation tool depicting the relative tradeoffs in true and false positives, plotting the true positive rate (recall, sensitivity) on the $y$ axis and the false positive rate on the $x$ axis [71]. It is common to reduce ROC curves to a single summarizing cross-model comparison metric: the area under the ROC curve (AUROC), which has the statistical property equivalent to "the probability that a randomly selected presence will have a higher continuous prediction than a randomly selected absence across all thresholds" [72] (see also [71] p. 868, [73]). The AUROC has been used extensively in SDM [19, 20, 44, 74–76], in part because it is threshold independent and in part because it is robust to class imbalance, which is prevalent in SDM data. Despite its ubiquity, AUROC is not without its criticisms [76–78], and its dependence on true absences may not be desirable in the context of presence-absence models with background sampling.

The location on the the ROC curve that jointly optimizes the true and false positive rates is captured by the True Skill Statistic [79] (TSS), variously termed the Pierce skill score (after [80]), Youden's J [81], and the Hanssen-Kuipers discriminant [82], per Liu *et al.* [83]. This location represents a Pareto front [84], and the maximum value of the index can be used to identify the threshold at which continuous probability distribution can be divided into binary classes to balance error types. The TSS has been studied in the context of SDM [19, 75, 85], and is recognized as an alternative to Cohen's kappa ([86], that avoids dependence on prevalence [75]. Leroy *et al.* [69] demonstrate that TSS actually does depend on prevalence in some contexts, though Allouch *et al.* [75] argue that with TSS, the effects of prevalence can be interpreted as evidence of real ecological phenomena, as opposed to statistical artifacts. For each of our predictions, we derived the cross validation mean AUROC and maximum TSS, along with the associated thresholds that maximized TSS. In that our industry

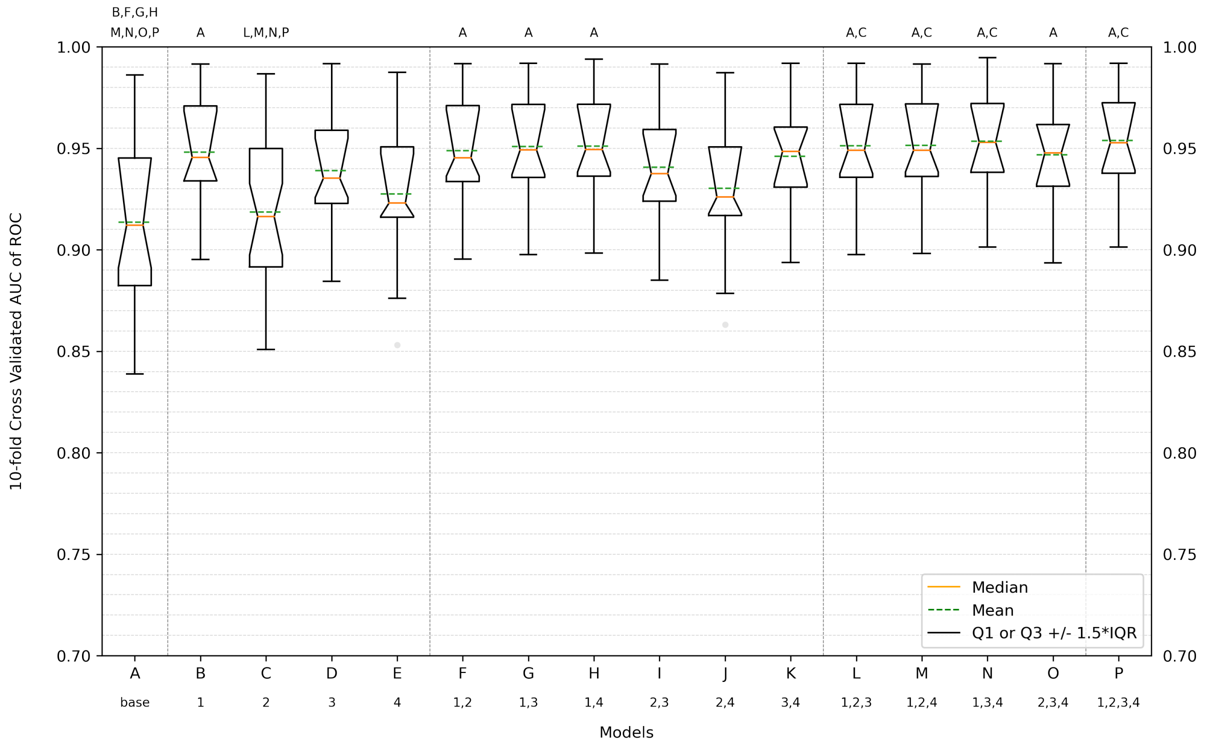


**Fig. S4**: Exploring explanatory variable selection. 10-fold cross validated AUC of ROC for auto-tuned MaxEnt models applied to each of 21 mills (each box) in a tuning subset and each of 16 different models (A-P). All models contain base terms, where base = costDistance, costDistanceFocalMean, costDelta12, and costDelta13 (see S1). Each subsequent model contains base plus combinations of: 1 = distanceToUrban, 2 = distanceToPalm, 3 = population density, and 4 = road density, listed below the x axis. Letters in the upper margin denote models for which there is a statistically significant difference ($\alpha = 0.05$). Boxes cover the first through third quantiles (hinges), whiskers of 1.5×interquartile range, and further outliers (points). Median and mean values are shown in green and orange, respectively.

standard model did not yield probabilistic outputs, its TSS depends on a single error matrix per cross validation fold.

###### *1.5.3.2 AUPRG and $F_1$*

Precision-recall curves are similar to ROC curves in providing threshold-independent measures of model performance, but consider the relationship between precision on the $y$ axis and recall (sensitivity, true positive rate) on the $x$ axis [72, 87]. Like ROC curve analysis, use of precision-recall curves is common in binary classification tasks, particularly when there is class imbalance or emphasis is consciously placed on the positive (minority) class, as neither precision nor recall depend on the false negatives [71, 87, 88]. Precision-recall curves have been employed in the context of SDM, and can help avoid measures of accuracy that depend on prevalence of both classes, as is the case with ROC metrics [72, 83, 88]. A number of studies have reported the area under the precision-recall curve as a single cross-model comparison metric [72, 87, 89, 90].

The analog to TSS when working with precision-recall curves, is the $F_\beta$ metric [91, 92], which has also been used in the context of SDM [93–95]. As with TSS, $F_\beta$ represents a joint optimization (balancing) of precision and recall, and the maximum value of the index can be used to identify the threshold at which continuous probability distribution can be divided into binary classes. Using the area under the recision-recall curve in combination with $F_\beta$ values is inherently problematic, as the area under the curve depends on an arithmetic mean of precision values, while the corresponding $F_\beta$ depends on the harmonic mean [84]. Flach and Kull's [84] precision gain and recall gain resolve this incompatibility, yielding curves that all share the desirable properties of ROC curves, permitting valid reporting of the area under the precision-recall gain curve (AUPRG) in conjunction with associated $F_\beta$ measures. Here, we derived the cross validation mean AUPRG and the maximum precision-recall gain-based $F_1$ accuracy for each of our predictions, along with the thresholds that maximized $F_1$ accuracy. As with TSS, the $F_1$ scores we report for our industry standard model depends on a single error matrix per cross validation fold.

###### *1.5.3.3 Kappa*

Cohen's kappa [86] is a widely used measure across numerous fields [96–98]; >43700 citations as of May 2022, per Google scholar), and seeks to correct overall accuracy by accounting for chance occurrence. In the context of SDM, the use of kappa has been criticized for its dependence on prevalence — it is generally not suitable when there is gross class imbalance [69, 72, 75, 99]. However, as with our other metrics, there are contradictory findings [100]. For each of our predictions, we derived the cross validation maximum kappa, along with the associated thresholds that maximized kappa. As with TSS and F1, the kappa scores we report for our industry standard models depend on a single error matrix per cross validation fold.

###### *1.5.3.4 Statistical comparison and prediction to grid*

We were interested in a single modeling framework that, on average, yielded superior predictive results across all 1,570 mills. For this reason, we compared our models using one-way analysis of variance (ANOVA; [101]), which was applied independently to each set of five distributions associated with the above-mentioned accuracy assessment metrics. Where omnibus $F$-tests indicated significant differences in accuracy between model types, pairwise comparisons were performed with Tukey honest significant difference tests [102]. Our top performing model was then used to produce continuous valued predicted probability surfaces for each mill, extending across the mills-specific prediction domains.

[10] Orbital Insights. Geolocation Traceability. URL https://docs.orbitalinsight.com/docs/

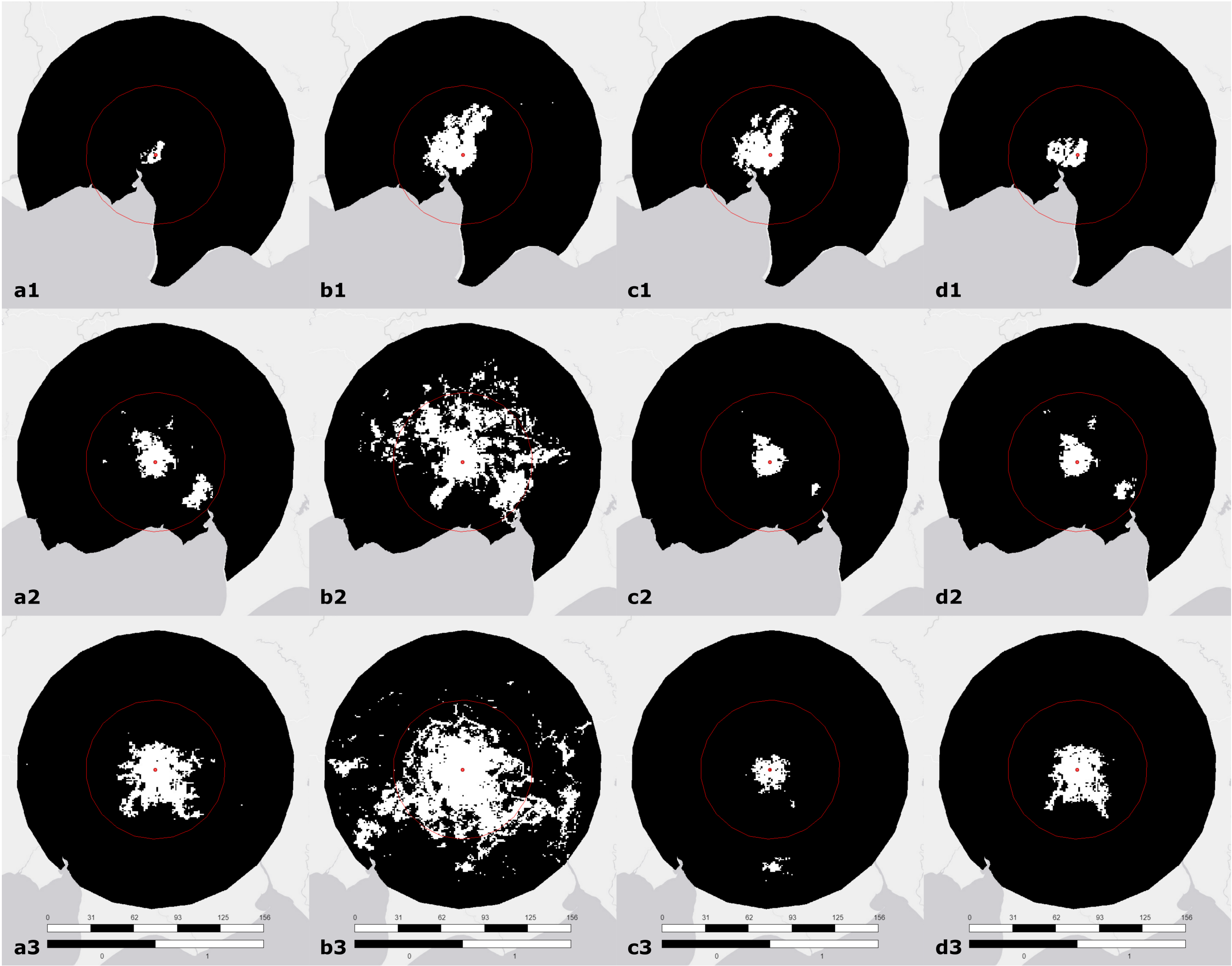


**Fig. S5**: Classified probability surfaces from main text Fig. 2, corresponding with for mills detailed in main text Table 1 (rows 1-3), for each of four models: MaxEnt (a), down-sampled random forests (b), down-sampled gradient boosted regression trees (c), and an ensemble (d). Classification was performed using the optimized precision-recall gain-based $F_1$ thresholds established for each mill (see main text Fig. 1 "Maximized PRG F1"). Red circular region corresponds with the current industry standard 50 km radial sourcing model. Red point marks mill location. Legend: upper bar is scale, in km; lower bar predicted class, where 1 = part of the focal mill's sourcing domain, and 0 = not part of the focal mill's sourcing domain.

geolocation-traceability.

[17] Franklin, J. *Mapping species distributions: spatial inference and prediction* (Cambridge University Press, 2010).

[45] Breiman, L., Friedman, J. H., Olshen, R. A. & Stone, C. J. *Classification and regression trees* (Routledge, 1984).

[52] James, G., Witen, D., Hastie, T. & Robert, T. *An Introduction to Statistical Learning* (Springer, 2017).

[64] Palm, F. C. & Zellner, A. To combine or not to combine? issues of combining forecasts. *Journal of Forecasting* **11** (8),

687–701 (1992). https://doi.org/10.1002/for.3980110806 .

[80] Pierce, C. The numerical measure of the success of predictions. *Science* **ns-4** (93), 453–454 (1884). https://doi.org/10.1126/science.ns-4.93.453.b .

[87] Davis, J. & Goadrich, M. *The Relationship between Precision-Recall and ROC Curves*,

233–240. ICML '06 (Association for Computing Machinery, New York, NY, USA, 2006). URL 10.1145/1143844.1143874.

[101] Weisberg, S. *Applied Linear Regression* Fourth edn. Wiley Series in Probability and Statistics (John Wiley & Sons, 2014).

[102] Tukey, J. W. *Exploratory data analysis* Vol. 2 (Addison, 1977). URL http://theta.edu.pl/wp-content/uploads/2012/10/exploratorydataanalysis_tukey.pdf.
